# Tumor-specific draining lymph node CD8 T cells orchestrate an anti-tumor response to neoadjuvant PD-1 immune checkpoint blockade

**DOI:** 10.1101/2025.04.27.650862

**Authors:** Rachel Honigsberg, Tatiana Cruz, Liron Yoffe, Meixian Stephanie Tang, Ozge Dicle, Geoffrey Markowitz, Marissa Michael, Arshdeep Singh, Nasser K. Altorki, Olivier Elemento, Jonathan Villena-Vargas

## Abstract

Elucidating the anti-tumor role of tumor-draining lymph nodes (tdLNs) in patients could offer critical mechanistic insight and shift therapeutic strategies from a tumor-centric approach to one that considers tumor-immune system interplay. Our study characterizes benign tdLNs T cell anti-tumor responses beyond initial T cell priming in patients with resectable non-small cell lung cancer. We further investigated whether tumor-specific tdLN T cells were altered by immune checkpoint blockade (ICB) locally and systemically. We performed single-cell TCR lineage tracing and transcriptomic profiling on 672,886 CD8 T cells from 41 tumor, benign tdLN, and blood samples in 14 patients treated with or without neoadjuvant chemoimmunotherapy (ChemoIO). Using deep-integrating clonal tracking with machine learning-based transcriptional analysis, our findings revealed that benign tdLNs locally and independently orchestrate two transcriptionally distinct tumor-specific memory CD8 T cell populations: one with ZNF683+ CXCR6+ tumor tissue-residency potential, and another with cytotoxic memory potential. Furthermore, tdLN-derived clones not only constitute the dominant tumor-infiltrating (75%) and circulating (>90%) tumor-specific expanded T cell populations but also preserve their transcriptionally distinct subset identities within the tumor T cell effector state. ChemoIO selectively increased the clonal diversity and cytotoxic memory/TEMRA programs of tdLN-derived clones locally and systemically, both of which remained unchanged in clones lacking tdLN TCR lineage. In conclusion, the tdLN locally orchestrates tumor-reactive and ChemoIO-reactive transcriptional distinct T cell subsets that shape the circulating blood and tumor T cell environments. These findings represent a clinical paradigm shift with implications regarding the extent of tdLN resection during surgery, timing of ChemoIO treatment, and the development of memory T cell-based immunotherapies.

## Main

Evidence-based changes to cancer treatment protocols are difficult to achieve. In particular, the anti-tumor role of tumor-draining lymph nodes (tdLNs) and the consequences of their surgical removal have been challenging to evaluate in patients. With the advent of single-cell analysis and deep T cell receptor (TCR) sequencing, it is now possible to investigate the origins and dynamics of T cells between tdLNs and tumors, and to assess how immunotherapy modulates these populations.^1,2^ This understanding may improve the success of chemoimmunotherapy (ChemoIO) and inform strategies to enhance patient outcomes.

Three key clinical questions are under active investigation in non-small cell lung cancer (NSCLC): (1) the appropriate extent of lymphadenectomy, (2) the optimal timing of ChemoIO (neoadjuvant vs. adjuvant), and (3) the identification of novel immunotherapy targets. Elucidating the role of benign tdLNs in anti-tumor immunity could provide critical mechanistic insight into these questions and a paradigm shift of therapeutic strategies from a focus on events within the tumor towards the interplay between the tumor and the systemic immune response.

First, lymphadenectomy is routinely performed, yet the optimal extent of nodal dissection remains undetermined.^3–5^ While some clinicians suggest lymph node removal may limit metastatic spread, others argue it may eliminate a compartment critical for systemic immune activation.^3–9^ The sustained contribution of tdLNs to anti-tumor T cell responses— beyond initial priming—remains uncertain in patients, leaving the immunological impact of their removal unclear.

A second major area of debate is the timing of ChemoIO, which remains divided between adjuvant and neoadjuvant approaches.^10–15^ Emerging data suggests that tdLNs may support effective immunotherapy responses.^9^ Preclinical studies show that both lymphadenectomy and blocking T cell migration from tdLN to the tumor using FTY720 reduce the efficacy of PD-1 checkpoint blockade.^9,16–18^ However, no patient studies have compared the phenotypes of tumor-specific tdLN T cells in ChemoIO-treated versus untreated individuals. Such data could justify adjusting treatment timing to preserve tdLN function before surgical resection.

Third, memory T cells have emerged as promising candidates for novel immunotherapies, yet strategies to effectively target them remain undeveloped.^18–20^ Memory T cells play a crucial role in maintaining systemic anti-tumor immunity, including repopulating exhausted cells and surveilling for metastatic disease.^19,21,22^ The tdLN harbors memory-like tumor-specific CD8 T cells capable of engaging in immunoregulatory PD-1/PD-L1 interactions with dendritic cells, positioning the tdLN as an unexplored potential site of therapeutic action during ChemoIO.^16,23,24^ Demonstrating that memory-like tdLN T cells respond locally to ChemoIO and drive systemic and tumor effects would redefine them as a novel, actionable therapeutic target.

Despite its promise, studying benign tdLNs has been limited by several factors: (1) difficulty in reliably identifying draining nodes, (2) the polyclonal nature of tumor-specific T cell responses, (3) a lack of flow cytometry markers distinguishing tumor-reactive from bystander T cells in tdLNs, (4) limited availability of patient samples, and (5) costs.^2,3,18,19,25–32^

Paired single-cell RNA and TCR sequencing now enables high-resolution tracking of tumor-specific T cells across tissues, surpassing the limitations of flow cytometry and tetramer-based analyses.^1,2,18,31^ To date, Pai et al. and Huang et al. have conducted the only patient studies examining systemic tumor-specific T cell responses within tdLNs using this technique.^2,18^ Pai et al. identified a progenitor exhausted memory subset in tdLNs from three PD-1 blockade–treated NSCLC patients.^2^ Huang et al. found stem-like tumor-specific tdLN CD8 T cells in six untreated hepatocellular carcinoma patients, which when transferred to a D-1/PD-L1–treated mouse model mediated superior responses.^18^ However, both studies lacked comparative analyses between treated and untreated patient cohorts and/or benign tdLN samples—key controls needed determining the role of tdLNs in therapeutic immune response in patients.

To address these limitations, we performed integrated single-cell transcriptomic and TCR lineage analysis on 672,996 CD8 T cells from 41 tumor, benign tdLN, and peripheral blood samples collected from 14 patients with resectable NSCLC treated with or without neoadjuvant ChemoIO. Using TCR lineage tracing, we further examined how tdLN-derived clones behaved in the tumor and blood compared to clones restricted to the tumor or the tumor and blood. In this paper we aimed to determine whether regional benign tdLNs are a central source of therapeutic T cell immunity in resectable NSCLC with or without the effect of ChemoIO.

## Results

### Tumor-specific clones in surgically resected NSCLC with distinct tissue lineages are found in the tdLN, blood, and tumor

To characterize the systemic CD8 T cell response following neoadjuvant chemotherapy combined with PD-1 ICB, we collected the tumor, tdLN (defined as regional intralobar N1 lymph nodes closest to the tumor) and peripheral blood from clinically matched early-stage NSCLC patients. Samples included tumor, tdLN, blood samples from eight patients who received neoadjuvant PD-1 ICB (Nivolumab or Pembrolizumab) and platinum-based chemotherapy and from six treatment naïve patients.^12^ Samples were then sorted for CD45+ CD3+ CD8+ T cells were sequenced for paired single cell (sc)RNA and scTCR (**Fig. 1a**). A T cell clone was defined by a collection of T cells with the same unique combination of all TCR alpha and TCR beta CDR3 amino acid regions. ^1,2,31,32^ From the 672,886 CD8 T cells sequenced, 390,375 CD8 T cells had both TCR alpha and TCR beta regions sequenced and scRNA sequencing (**Fig. 1b**).

**Fig. 1:**
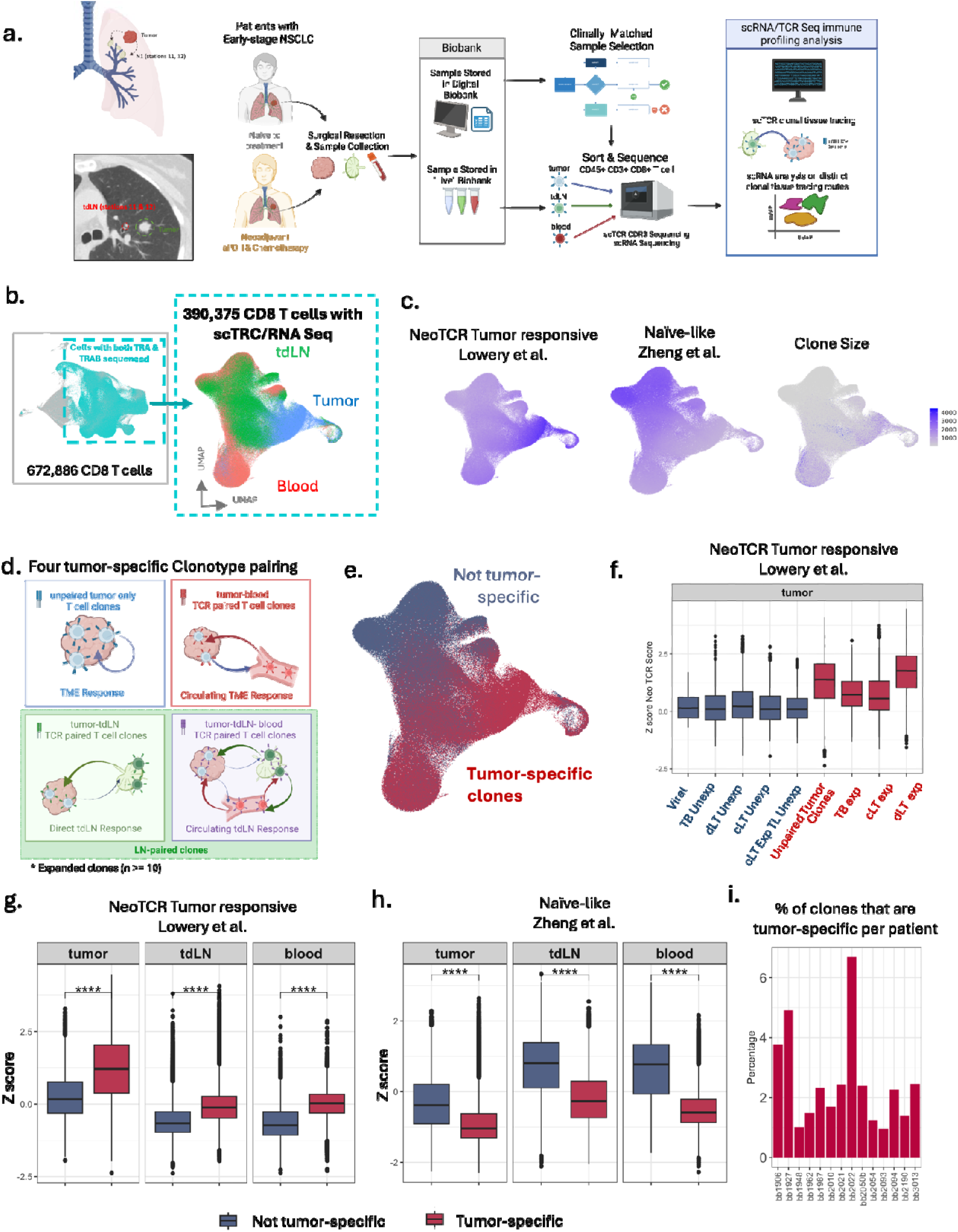
Defining and validating tumor-specific T cell clones in early-stage non-small cell lung cancer. a. Schematic of patient sample collection sorting, sequencing and analysis pipeline. b. Selection of sequenced CD8 T cells (672,886 total) which have both scRNA and alpha and beta chains TCR sequenced (390,375 total) followed by dimensional UMAP dimensional reduction. c. Feature plots of NeoTCR tumor responsive signature, naïve-like signature and clone size.^32,34^ d. Schematic of tumor-specific clone-type pairings based on TCR tissue lineage tracing. e. UMAP labeling T cell tumor-specificity based TCR tissue lineage tracing described in d. f. Validation of tumor-specific clone type pairing categorization. Clonal average z-score of tumor T cell NeoTCR tumor responsive signature by TCR tissue lineage tracing, expansion and if the TCR was found as a viral epitope on the VDJ database. g. NeoTCR tumor responsive signature (Lowery et al.) clonal average z-score applied to each tissue by clonal tumor-specificity based on TCR lineage tracing described in d. (t-test, p < 0.0001).^32^ h. Naïve-like signature (Zheng et al.) clonal average z-score applied to each tissue by clonal tumor-specificity based on TCR lineage tracing described in d. (t-test, p < 0.0001).^34^ i. Percent of paired sc RNA/TCR clones that are tumor-specific per patient.

Clonal tissue lineage tracing was performed using the TCR CDR3 region as a clonal barcode across tissues. Clones were then categorized based on the presence of their TCR sequences in a viral epitope in a TCR VDJ database, their presence across tissues, and whether the clones were expanded (≥10 T cells) or unexpanded (< 10 T cells). Previous studies have described clonal expansion within the tumor relating to tumor-reactive T cells. ^1,2,31,32^ Therefore, we defined clonal tumor-specificity as expanded clones that have T cells present in the tumor. Potential “bystander” CD8 T cells defined by TCRs listed as viral epitope species in a TCR VDJ database and unexpanded clones were not included as tumor-specific clones (**Fig. 1d**).^1,2,31–33^

Tumor-specific clonotype categories included unpaired tumor-only clones (uT) with expanded TCRs only found in the tumor, LN-paired expanded clones (tumor-specific clones with TCR lineage with the tdLN), and tumor-blood clones with TCR only present in tumor and blood (bT). LN-paired clones were subdivided into two categories: circulating LN-paired clones (cLT) found in tumor and tdLN and circulating in peripheral blood, and direct LN-paired clones (dLT) that had TCR lineage only between tdLN directly to tumor (**Fig. 1d**). Of the 112,553 unique TCR clones found across all three tissues, 4% of clones were tumor-specific, on average 2.5% of clones per patient (**Fig. 1i**).

This clonal labeling was then validated by comparing non-tumor-specific clones (viral, unexpanded, not clonally linked to the tumor) and tumor-specific T cell clonal NeoTCR CD8 T cell tumor-reactivity score from Lowery et al. and Naive-like score from Zheng et al.^32,34^ Within the tumor, tdLN and blood, tumor-specific clone categories were found to increase in tumor-reactivity scoring and decrease in Naive-like scoring relative to non-tumor-specific clone categories. Interestingly, viral clones had low NeoTCR tumor-reactivity but were less naive than the unexpanded clones (**Fig. 1f-h**).^32,34^ Circulating LN-paired clones that had less than 10 tumor and tdLN CD8 T cells had a low NeoTCR score and higher Naive score, potentially representing another subset of bystander T cells. ^32,34^ Therefore, circulating LN-paired clones were only considered tumor-specific if there were at least 10 cells found within the tumor and tdLN. (**Fig. 1f**).

### Tumor-specific clonal profile across tdLN, blood, and tumor

Prior scTCR/RNA clonal analysis in patients of tdLNs have centered on metastatic disease. ^2^ We instead examined how the tissue environment shapes tumor-specific CD8 T cells of early-stage NSCLC patients. We compared functional scores and gene expression profiles of tumor-specific clones within the blood, tdLN, and the tumor. Tumor-infiltrating T cells were the most effector-like, terminally exhausted, with high expression of CD39+CD69+ terminal CD8 T cells signatures.^34,35^ In contrast, tdLN and blood T cells had higher Naive/CM-like and memory-like scores. (**Supplement Fig. 1**)^34,35^ Circulating T cells exhibited increased TEMRA and cytotoxic signatures relative to tdLN, consistent with prior reports.^34,35^ Tumor T cells also had elevated expression of TEMRA compared to tdLN.^34^ TdLN T cells expressed the memory marker *IL7R*, whereas blood T cells expressed *CX3CR1* and *KLF2*. (**Supplement Fig. 1**). Within both dLT clones and cLT clones, tdLN T cells displayed increased Naive-like scores, compared to their tumor counterparts, which expressed tissue resident memory and dysfunction markers (**Supplement Fig. 1**).^34,35^ We hypothesize that, in the absence of chronic antigen and immune suppression, CD8 T cells remain memory-like within the benign tdLN and become dysfunctional upon entry into the TME.

### Clonal lineages from tumor-draining lymph nodes shape the intratumoral CD8 T cell landscape

Prior literature has established the TME as central to T cell differentiation and exhaustion.^18,20,36^, However, we aim to establish if tdLN lineage was also a contributing factor to the tumor T cell phenotype. To do this, we compared uT clones restricted to the tumor and LN-paired clones in the tumor. uT clones represented the impact of the TME, without the contribution of the tdLN. On the other hand, LN-paired (dLT and cLT) clones represented the contribution of the tdLN in the tumor T cell response.

While it has been established that tdLN-activated T cells migrate to the tumor, the capacity of the tumor that comprises these tdLN-derived T cells has yet to be defined.^36,37^ The proportion of tdLN TCR-linked cells within the tumor was higher than clones only found in the tumor. On average, tdLN-linked cells accounted for 75% of tumor T cells per patient, while uT clones contained 22% of tumor T cells, and bT clones were only 3% of cells. (**Fig. 2b**) Tumor-specific clones paired to the tdLN were more clonally diverse than uT clones--measured as the number of unique TCRs per 5000 cells per patient. On average, there were 73.6 tumor-specific clones with tdLN lineage per 5000 cells per patient, compared to 22.9 uT clones (**Fig. 2a**). Previous studies have proposed that clonal diversity may enhance the ability of T cells to recognize tumor antigens and drive a more effective anti-tumor response.^37^ These findings suggest that the tdLNs have a significant contribution towards the clonal landscape of tumor-infiltrating T cells.

**Fig. 2:**
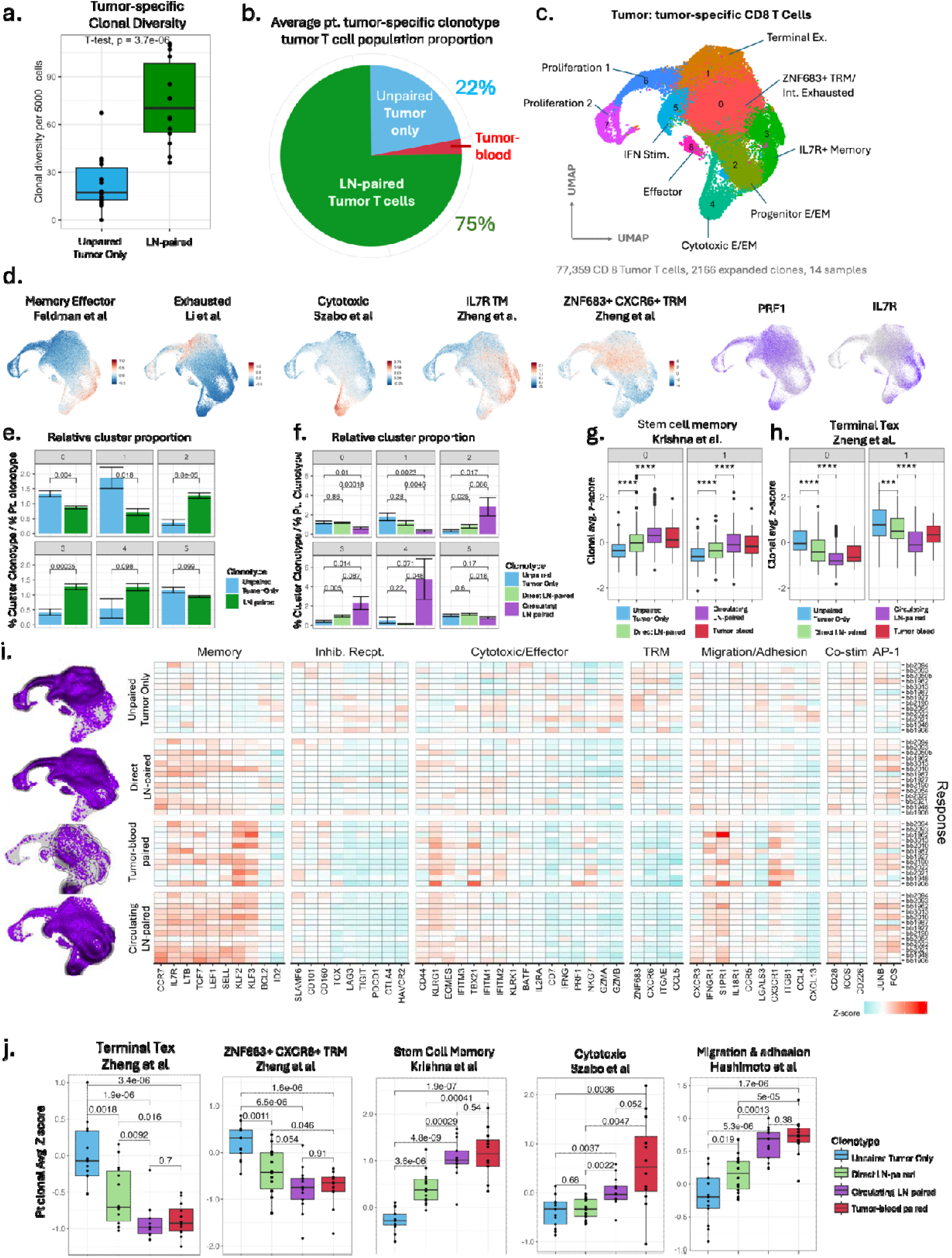
LN-paired T cells within the tumor are distinct from unpaired tumor T cells. a. Patient tumor-specific clonal diversity (number of unique TCRs per 5,000 cells) of unpaired tumor only clones versus LN-paired clones (paired t-test, p < 0.0001). b. Average patient tumor-specific tumor T cell population proportion based on clonal lineage with the LN (clone type pairing described in 2.1 d.). c. UMAP dimensional reduction of tumor-specific CD8 T cells, including 77,359 T cells and 2,166 expanded clones, 14 patient tumor samples. Nine T cell cluster subsets were generated. d. Feature plot of signatures relating to T cell function normalized using z-score (mean = 0, STD = 1) and of gene expressions *PRF1* and *IL7R*.^31,34,38,39^ e. Relative clone type cluster proportion of unpaired tumor only clones and LN-paired clones (percent of clone type T cells within cluster for each patient/percent of clone type T cells for each patient). A value of 1 represents a cluster with clone type population proportions the same as the patient tumor clone type population proportions. f. Relative clone type cluster proportion of unpaired tumor only clones, direct LN-paired clones, and circulating LN-paired (percent of clone type T cells within cluster for each patient/percent of clone type T cells for each patient). A value of 1 represents a cluster with clone type population proportions the same as the patient tumor clone type population proportions. g-h. The clonal average z-score of (g.) stem cell-like and (h.) terminal exhaustion signatures within ZNF683+ TRM/Int Exhausted (cluster 1) and Terminal Ex. (cluster 0) clusters by clone type pairing.^34,35^ i. Heatmap of average z-score clonal expression levels per patient of genes representing a range of T cell functions (memory, inhibitor receptors, cytotoxic/effector, etc.) for each tumor-specific clonal type pairing category (unpaired tumor only clones, direct tumor-tdLN clones, circulating LN-paired clones, tumor-blood). The left of the heatmap are the density plots of cells per clone type pairing. j. Patient clonal average z-score of signatures from literature relating to T cell function by clone type within the tumor (t-test, ANOVA, all p < 0.0001, except ZNF683+ CXCR6+ TRM p < 0.01).^34,35,38,42^

We next explored the tumor T cell cellular landscape through clonal transcriptomic analysis using unsupervised dimensionality reduction of all 77,349 tumor-specific CD8 T cells (2,166 expanded clones) from 14 patients (**Fig. 2c**). Nine distinct clusters and T cell subtypes ranged from memory state to dysfunctional and exhausted. Three clusters were more memory-like (IL7R+ Memory, Progenitor E/EM, and cytotoxic E/EM), defined by an increase in memory signatures from the literature and markers *IL7R*, *TCF7*, *LEF1*, and *SELL.*^1,2,31,32,34,35,38–42^ IL7R+ Memory was defined by increased expression of *IL7R* and low expression of activated, cytotoxic, and effector genes, while the Progenitor E/EM and Cytotoxic E/EM clusters showed evidence of effector activation. Progenitor E/EM and cytotoxic E/EM memory functions differed in progenitor capacity (*TOX, EOMES, GZMK, CD44*), cytotoxicity and migration (*KLF2, KLF3, CXCR1*). ^1,2,31,32,34,35,38–42^ Four clusters represented a variety of differentiated T cells (Effector, ZNF683+ Tissue Resident Memory Intermediate Exhausted, IFN stimulated, and Terminally Exhausted), defined by upregulation of effector (*PRF1*, *CD44*, *GZMB*, etc.) and inhibitory receptors (*HAVCR2, TIGIT, CTLA4*) but low expression of memory markers. ^1,2,31,32,34,35,38–42^ While all of these clusters have some expression of tissue resident memory (TRM) genes, the Intermediate Exhausted cluster had the highest expression of *CXCR6* and *ZNF683* and the highest z-score of Zheng et al.’s ZNF683+ CXCR6+ TRM signature.^34^ In contrast, the Terminally Exhausted cluster displayed the highest expression of inhibitory markers and signatures relating to exhausted (Li et al.), terminal exhaustion (Zheng et al.) and Exhausted TRM (Pauken et al.).^31,34,38^ IFN-stimulated T cells had increased genes relating to T cells responding to IFN stimulation (*STAT1, STAT2, MX1*).^31,32,34,35,38,39^ Lastly, we determined two clusters of proliferation, defined by the expression of *MKI67* (**Fig. 2d, Supplement Fig. 2**).^2,31,39^

To assess the phenotypic influence and contribution of tdLN clonal lineage within the tumor, we compared the relative abundance of LN-paired and uT clones across UMAP-defined clusters. For each patient, we calculated the proportion of LN-paired and uT clones within each cluster, relative to that patient’s global clonotype proportion. A relative abundance of 1 would relay a similar cluster clonotype proportion to that of the patient, and no influence of clonotype (LN-paired or uT) on that cluster. Notably, tdLN clonal lineage favored the memory-like clusters IL7R+ Memory and Progenitor Exhausted, while uT clones had a larger contribution within the most exhausted clusters (Int exhausted TRM and Terminally Exhausted). These findings indicate that the tdLN preferentially contributes effector memory and progenitor memory-like clones to the tumor (**Fig. 2e**).

### Two transcriptionally distinct LN-paired clones are found within the tumor: directly LN-paired (dLT) and circulating LN-paired (cLT)

We next investigated whether there were differences between the dLT and cLT clones within the tumor. First, we compared the relative cluster abundance of dLT clones and cLT clones. The dLT clones had an increased proportion in the Terminally Exhausted subset and the Intermediate TRM cluster subset, while the circulating LN-paired clones had high contribution on the Cytotoxic E/EM cluster. This displays that dLT clones have a closer differentiation resemblance to uT clones compared to the more cytotoxic profile of cLT clones (**Fig. 2f**).^34,35,38^

We extended our analysis with a more granular comparison of average clonal expression levels per patient, focusing on genes representing a variety of T cell functions for each clonotype pairing category (uT, dLT, cLT, bT), followed by further analysis by applying signatures from the literature.

As expected, all four clonotype pairings within the tumor had high expression of genes relating to cytotoxic/effector function. However, cytotoxic/effector genes were specific to certain clonal pairings, indicating potentially different effector functions within the tumor, suggesting functional heterogeneity. bT clones had heightened expression of *KLRG1, NKG7* and *TBX21* (T-bet).^40^ LN-paired (dLT and cLT) clones had an increased expression of *CD44* and *EOMES*. uT clones exhibited high expression averages of genes relating to exhaustion (e.g. *TOX, LAG3*) and tissue residency (*CXCR6, ZNF683, ITGAE,* etc.), but low expression of memory-like genes, except for *BCL2* and *ID2*. On the other hand, paired closed (dLT, cLT, bT) had high expression of stem cell memory and low expression of Tex and TRM signatures compared to uT clones. AP-1 activation markers *JUNB* and *FOS* were increased in paired clones and decreased in uT clones. ^34,35^ LN-paired clones had higher expression of *CD44, EOMES*, *CXCR3* and *CD28* compared to bT clones, while bT clones uniquely displayed an increase in *CXCR1*. These results indicate that although all clones within the tumor are exposed to the same TME, tdLN circulating blood clonal lineages impact clonal cell fate and may drive variable anti-tumor functions (**Fig. 2i**).

However, there were also transcriptional distinctions between LN-paired dLT and cLT clones within the tumor. cLT clones had high expression of migration and adhesion signatures within the tumor, specifically, *IFNGR1* and *S1PR1* were highly expressed (**Fig. 2i-j**). ^31,42^ Additionally, cLT clones had a higher expression of Li et al.’s cytotoxic vs. dysfunction signature compared to dLT clones (**Supplement Fig. 2**).^31^ While all paired clones exhibited higher expression of memory genes (*CCR7, IL7R, TCF7, SELL,* etc.), cLT and bT clones had an increase in *KLF2* and *KLF3* expression, which was not increased in dLT clones. ^40,43^ dLT clones on the other hand had a higher expression of TRM and Tex signatures compared to cLT clones (**Fig. 2i-j**).^34^ These results suggest that there are two distinct LN-paired anti-tumor clonal responses within the tumor: a cytotoxic migratory response from cLT clones, and a more activated TRM-like response from dLT clones.

We further examined transcriptional differences between dLT clones and uT clones within the terminally exhausted cluster to determine if LN clonal lineage contributed to differences within the most exhausted T cells. Terminally exhausted uT clones displayed higher expression of T cell dysfunction signatures (Li et al) while dLT clones had relatively higher expression of effector memory function (**Fig. 2g**, **Fig. 2h**).^38^ We hypothesize that this preserved memory plasticity may signify an effector subset previously implicated in LN-paired derived clonal replacement of exhausted dLT clones (**Fig. 2j, Supplement Fig. 2**).^19^

The clear differences between unpaired uT (T cells found exclusively within tumors) and paired clones (T cells clonally related to lymph nodes or blood) demonstrate that sc TCR sequencing effectively distinguishes these two populations. Notably, the cLT clones and bT clones show distinct cytotoxic functional profiles, suggesting that peripheral circulation may modify clonal fate within tumors. Although dLT (T cells likely migrating directly from LN into tumors) differ from uT clones, they do share similar functional traits, such as increased activation and exhaustion. Together, this indicates that dLT clones and cLT clones represent distinct subsets within tumors, each likely playing distinct roles in shaping anti-tumor immunity.

### LN-paired clones reside in an earlier cell state in the tdLN, mirrored in their tumor counterpart

TdLNs have been proposed as reservoirs of tumor-specific T cells.^9,44^ To determine whether the tdLN was a site for early differentiation, we characterized 51,249 tumor-specific tdLN CD8 T cells that belonged to 1,689 expanded clones. Unsupervised dimensionality reduction revealed 10 distinct memory and effector memory clusters. (**Fig. 3e**). All tdLN CD8 T cell clusters had high expression of memory genes (*TCF7, LEF1, IL7R*, etc.) (**Fig. 3f, Supplement Fig. 3**).^2,31,34^ These findings align with Pai et al., who demonstrated that tumor-specific tdLN T cells are generally progenitor-like.^2^ Cluster profiles ranged from more quiescent Naive/central memory (CM)-like, Early TRM, Cytotoxic Memory, Progenitor Exhausted, and to more differentiated Effector/Effector Memory function.

**Fig. 3:**
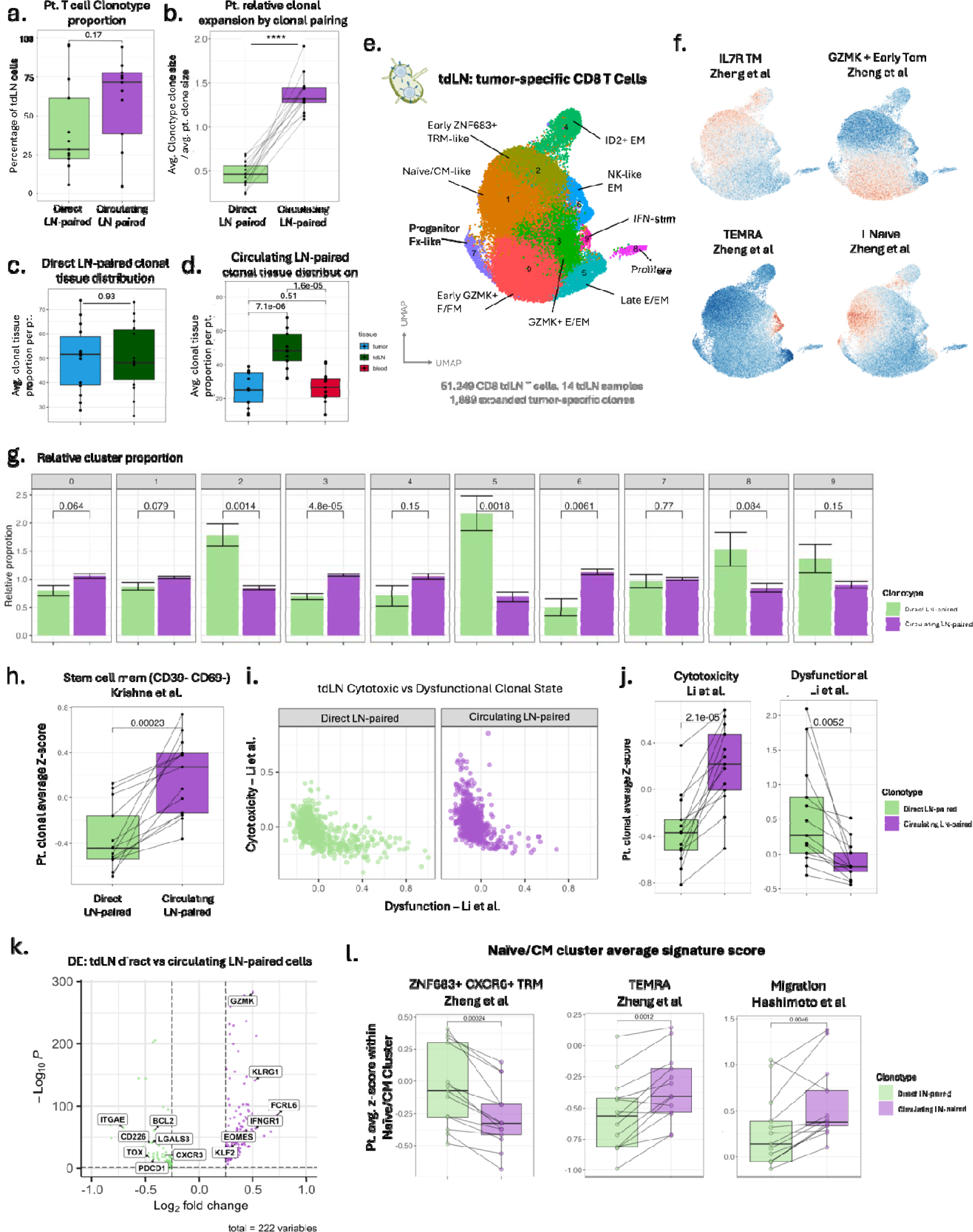
Clonal differentiation of LN-paired clones begins in the tdLN and reflects the subtypes seen in the tumor. a. Patient T cell clone type proportion within tdLN (paired t-test, p = 0.17). b. Clone type relative clonal expansion (average clone size) per patient (paired t-test, p < 0.0001). c-d. Average tissue proportion per clone per patient for direct LN-paired clones (c., paired t-test, p=0.93) and circulating LN-paired clones (d., paired t-test, ANOVA p < 0.0001). e. Dimensional reduction using UMAP of 51,249 tumor-specific CD8 tdLN T cells, including 1,689 expanded LN-paired tumor-specific clones from 14 patient samples. f. Feature plots of memory-like T cell signatures described by Zheng et al.^34^ g. Relative clone type cluster proportion of direct and circulating LN-paired T cells (percent of clone type T cells within cluster for each patient/percent of clone type T cells for each patient). A value of 1 represents a cluster with clone type population proportions the same as the patient tdLN clone type population proportions.^35^ h. Stem cell memory CD8 T cell average clonal signature z-score by clone type. i. Average clonal dysfunction versus average clonal cytotoxicity by Li et al. to distinguish cytotoxic and dysfunctional clones.^38^ j. Patient clonal average z-score of Li et al.’s cytotoxicity and dysfunction signatures.^38^ k. Differential expression between tdLN direct versus circulating LN-paired cells. l. Patient clonal average z-score signature expressions of T cells found within tdLN Naïve/CM cluster.

Although cLT clones comprised the majority (81.5%) of tdLN tumor-specific T cells, the clonal diversity between dLT clones and cLT was comparable on a per patient basis (**Fig. 3a**, **Fig. 3b**). dLT clones had balanced tissue distribution of T cells across tdLN and tumor tissues per patient, whereas cLT clones contained a higher proportion of T cells within the tdLN than the blood and tumor. Despite these differences in distribution, the number of cLT and dLT T cells within the tumor was similar in each patient (**Supplement Fig. 1**). These data suggest that cLT clones undergo preferential expansion within the tdLN, distinct from their tumor-restricted counterpart, highlighting a clonal expansion pattern unique to the lymph node compartment.

We next asked whether the memory-like tdLN environment shaped transcriptionally distinct dLT and cLT clones. Within the tdLN UMAP clustering, dLT clones were relatively abundant within the Early TRM and Late E/EM clusters, while cLT clones were expanded within the GZMK+ E/EM and cytotoxic NK-like memory clusters (**Fig. 3g**). Furthermore, dLT clones had higher expression of *CXCR6,* a gene relating to tissue residency and long-term survival in other cancers.^17,34^ On the other hand, cLT clones had increased *S1PR1* expression, potentially useful in peripheral migration and adhesion.^42^

Further differential expression analysis between direct and circulating LN-paired tdLN T cells revealed dLT clones were enriched for TRM, tumor-infiltrating, tissue-residency, activation, and progenitor exhausted-associated genes (*ITGAE, S100A11, CXCR4, PDCD1, TOX*), while cLT clones showed enrichment in cytotoxic/NK-like genes (*NKG7, KLRG1, FCRL6, KLF2*) (**Fig. 3e**).^2,40,45^ These differences were validated by comparing functional signature scoring from the literature. dLT clones also had an increase in ZNF683+ CXCR6+ TRM scoring and activation (**Supplement Fig. 3**).^34,38^ TdLN cLT clones exhibited greater cytotoxicity, elevated migration and adhesion signature expression, and enrichment for CD39-CD69-stem cell memory (**Fig. 3h**, **Fig. 3d, Supplement Fig. 3**).^35,38,39^ These results reveal a transcriptional imprint within the tdLN that is preserved across tissue clonal lineage and manifests as distinct differentiated differences in the tumor and peripheral blood.

Moreover, these tumor-relevant clonal differences in the tdLN were set in an early state of differentiation and memory. The cytotoxic markers that were found to be enriched in the cLT clones had increased expression in Naive/CM-like compared to dLT clones (**Supplement Fig. 3**). On the other hand, dLT clones were increased in expression of *ZNF684, CXCR6,* and *ITGAE* within the Naive/CM-like cluster, genes that are found in TRM tumor subsets, resulting in an increased expression of ZNF683+ CXCR6+ TRM T cell signature from Zheng et al (**Fig. 3f**).^34^ This suggests a differentiated separation of dLT and cLT clones starting at the earliest memory state of tdLN CD8 T cells that reflects the continued TRM and cytotoxic differences when found within the tumor.^35,38,39^

To further assess clonal divergence between cLT and dLT clones within the tdLN, we quantified the average clonal expression of cytotoxic and dysfunctional gene signatures defined by Li et al. to describe the two independent T cell differentiation trajectories.^38^ Based on these scores, each clone was categorized as cytotoxic (high cytotoxicity, low dysfunction), dysfunctional (high dysfunction, low cytotoxicity), or transitional—a pre-dysfunctional state along the exhaustion trajectory as previously described. dLT clones were predominantly classified within the transitional state, whereas approximately half of cLT clones exhibited a cytotoxic profile.^31^ Overall, cLT clones demonstrated elevated cytotoxicity and reduced dysfunction scores relative to dLT clones, indicating distinct differentiation trajectories shaped by lineage origin (**Fig. 3i, j; Supplement Fig. 3**).

### Tumor-specific blood CD8 T cells with tdLN lineage are more memory-like

To investigate the influence of tdLN clonal lineage on circulating tumor-specific CD8 T cells, we performed dimensionality reduction on 62,863 tumor-specific T cells, encompassing 1,156 expanded tumor-specific clones sequenced from peripheral blood. (**Fig. 4a**) This analysis identified seven transcriptionally distinct clusters, with the majority of cells partitioned into two dominant groups: memory (Naive/CM and *GZMK+* CM), and cytolytic TEMRA (cytotoxic E/EM, Super KILR E/EM, and effector E/EM). Memory clusters were enriched for canonical stem-like and homeostatic memory genes, including *TCF7, LEF1, IL7R, SELL.*^2,31,34^ Cytolytic clusters had lower expression of memory-like genes and higher expression of genes associated with NK-like cytotoxicity (*NKG7, IFITM1, IFITM2, IFITM3, FCGR3A,* etc.).^34,38,45–47^ Other clusters included a non-cytolytic CCR7-CD27+ CXCR1-population and a proliferative T cell cluster.^46^ Most clusters displayed expression of *TOX* and *TIGIT,* but low expression of *CTLA4* and *PDCD1* (**Fig. 4a**, **Fig. 4b, Supplement Fig. 4**).

**Fig. 4:**
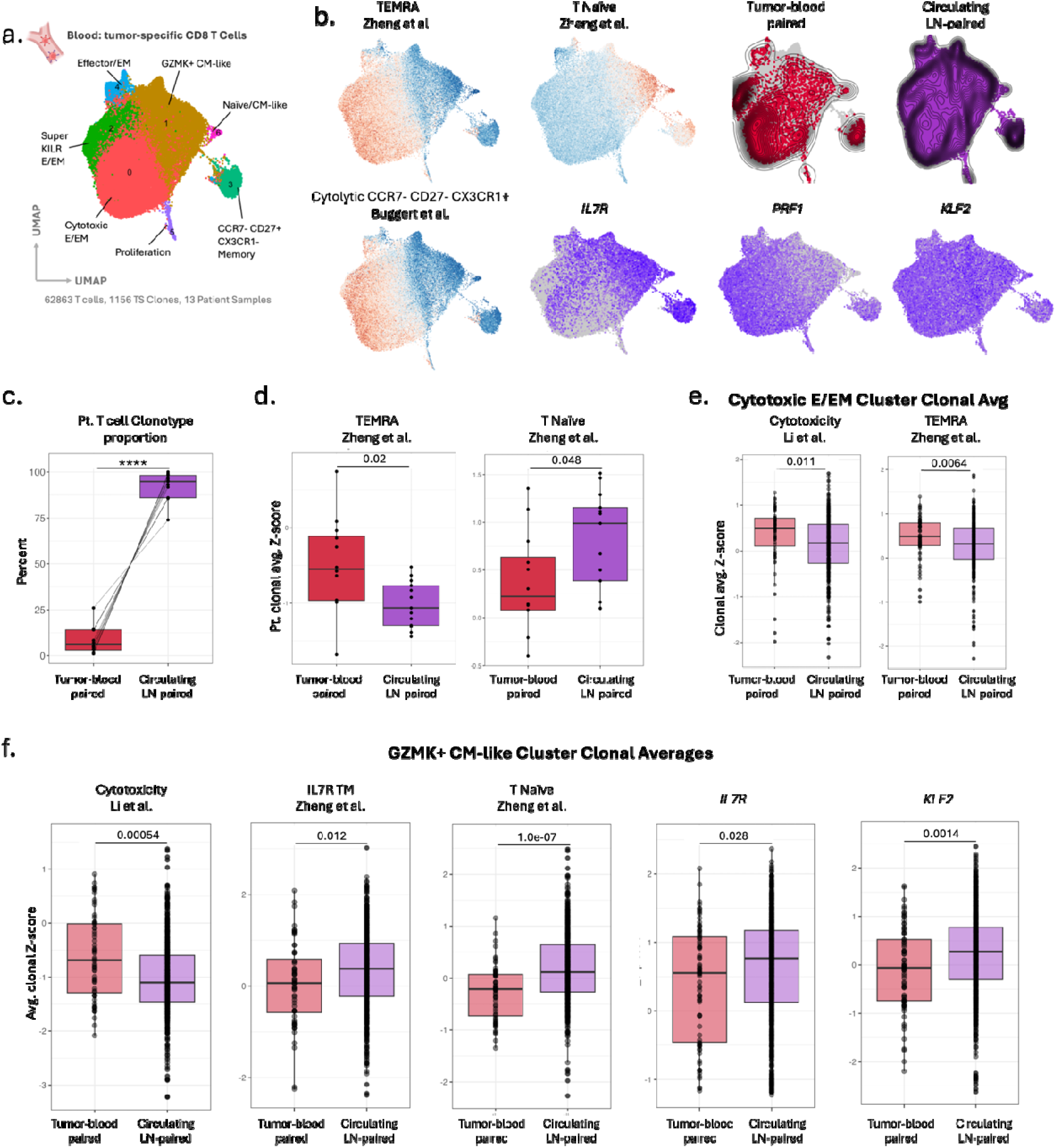
Contribution of LN-paired clones in circulating blood tumor-specific T cells. a. Dimensional reduction using UMAP of 62,863 tumor-specific blood T cells including 1,156 clones and 13 patient samples. b. Feature plots of T cell signatures from the literature and gene expression signatures (*IL7R*, *PRF1*, and *KLF2*).^34,46^ Top right are Dimplots highlighting tumor-blood paired and circulating LN-paired T cell UMAP locations overlaid by a density plot. c. Patient T cell clone type proportion within blood (paired t-test, p <0.001). d. Patient average clonal z-score of TEMRA (t-test, p=0.02) and T Naïve (t-test, p=0.048) signatures by Zheng et al.^34^ e. Patient clonal average z-score signature expressions of T cells found within the blood Cytotoxic E/EM cluster (t-test, p < 0.01).^34,38^ f. Patient clonal average z-score signature expressions of T cells found within the blood GZMK+ CM-like cluster (t-test, p < 0.05).^34,38^

Remarkably, an average of 95% of circulating blood tumor-specific CD8 T cells per patient were clonally linked to the tdLN (**Fig. 4c**). While both cLT clones and bT clones displayed cytotoxicity, cLT clones were consistently less differentiated (**Fig. 4d**). To further resolve phenotypic differences, we compared transcriptional signature differences within the Cytolytic E/EM and GZMK+ CM clusters, the largest memory and cytolytic clusters. Within the cytotoxic E/EM cluster, bT clones had increased expression of cytotoxicity and TEMRA-associated signatures.^34,38^ Similarly, within the GZMK+ CM cluster, bT clones had increased expression of the cytotoxic signatures, whereas cLT clones exhibited elevated memory-associated programs, including Naive-like and IL7R memory signatures (**Fig. 4e**, **Fig. 4f, Supplement Fig. 4**).^34,35^ These findings indicate that circulating tumor-relevant T cells clonally linked to the tdLN are less differentiated and retain a more memory-like transcriptional profile than those only linked to the tumor. This suggests that tdLN lineage may support the generation of circulating T cells poised for persistence.

### ChemoIO makes tumor-restricted clones less dysfunctional

To assess the impact of treatment on tumor-specific T cells across tissues, we compared the differences between patients treated with ChemoIO (n=8) and treatment-naïve patients (n=6). In the cohort of treated patients, there is an even distribution of non-responders (residual viable tumor [RVT] >30%, n=4), major pathological responders (RVT < 10%, n=3), and one partial responder (RVT = 30%, n=1) (**Fig. 5a**). In uT clones, ChemoIO reduced signature expression of dysfunction, terminal differentiation, terminal exhaustion, IFN stimulation and genes relating to proinflammatory cytokines, in line with prior literature describing a TIL ChemoIO response (**Supplementary Fig. 7**).^15,31,34,38,39^ Despite their differentiated state, these clones in treated patients retained greater memory potential, as evidenced by increased IL7R tissue-memory (TM) signature expression and elevated *IL7R* transcript levels (**Supplementary Fig. 7**).^34^

**Fig. 5:**
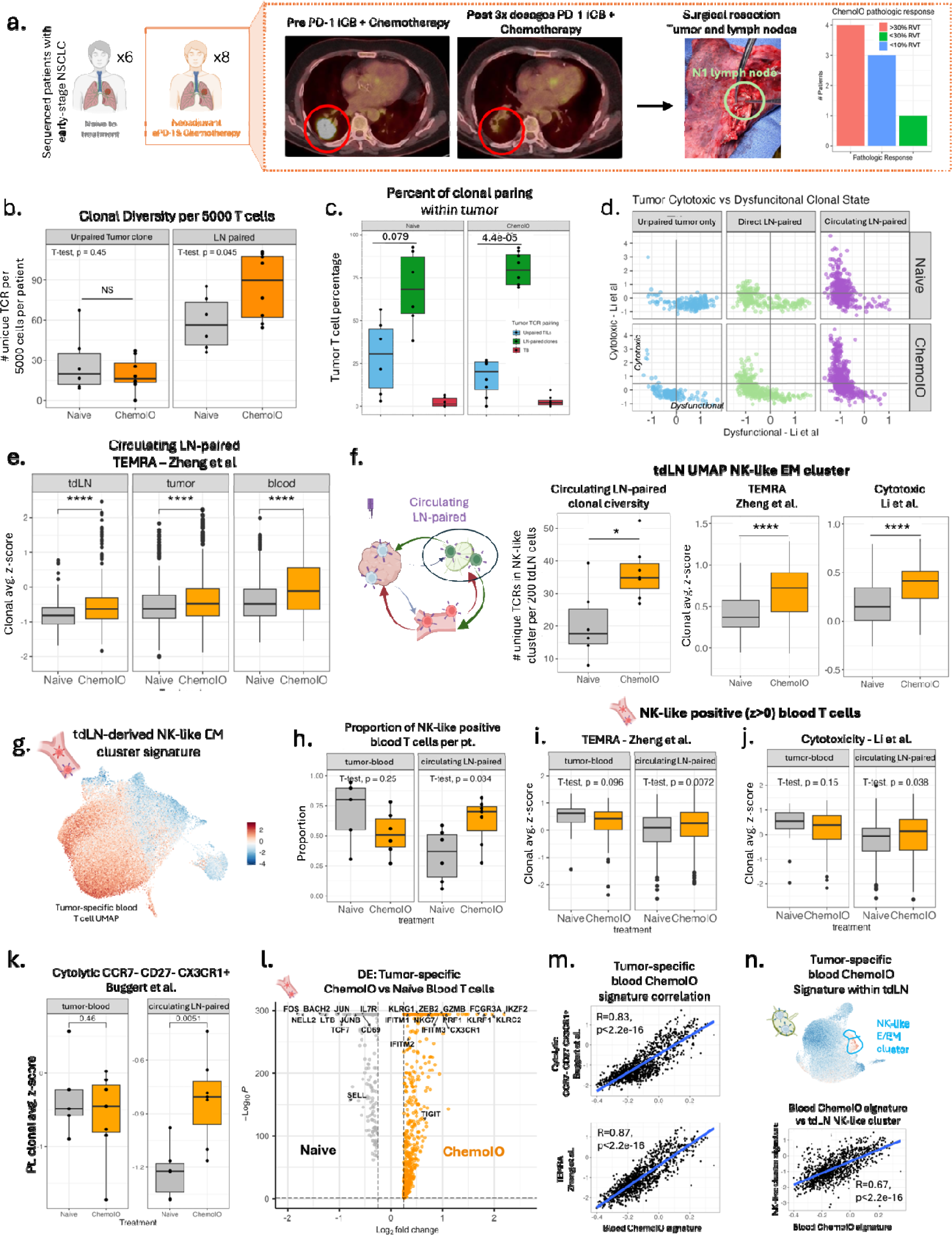
ChemoIO tumor-specific LN-paired response. a. CT scan of patient pre- and post-3 dosages of PD-1 ICB + chemotherapy followed by surgical resection of tumor and LNs. To the right is the pathologic response distribution of the eight patients treated by ChemoIO. b. Patient tumor-specific clonal diversity (number of unique TCRs per 5,000 cells) between treatment Naïve and ChemoIO treated patients for unpaired tumor only clones (t-test, p = 0.45) and LN-paired clones (t-test, p = 0.045). c. Tumor-specific tumor T cell population proportion based on clonal lineage with the LN (clone type pairing described in 2.1 d.) per patient (paired t-test between unpaired tumor only clones and LN-paired clones) by treatment. d. Average clonal dysfunction versus average clonal cytotoxicity by Li et al. to distinguish cytotoxic and dysfunctional clones separated by treatment and clone type pairing. Gating Cytotoxic clones to have > 0.5 average cytotoxic signature and < -0.25 average dysfunctional signature and Dysfunctional clones < 0.5 average cytotoxic signature and > -0.25 average dysfunctional signature.^38^ e. Treatment differences of average clonal z-score of TEMRA expression within circulating LN-paired clones within the tdLN, tumor and blood (t-test, p < 0.0001).^34^ f. Treatment differences in circulating LN-paired clones found within tdLN NK-like cluster: clonal diversity per 200 cells (t-test, p=0.05), clonal average TEMRA signature expression (t-test, p < 0.0001), and clonal average cytotoxicity signature expression (t-test, p < 0.0001).^34,38^ g. Feature plot on tumor-specific blood T cell UMAP of tdLN NK-like cluster signature (z-score normalized). h-j. Comparison of treatment response of blood T cells with tdLN NK-like positive (NK-like cluster signature high, z>0) within tumor-blood clones and circulating and LN-paired clones. h. Proportion of T cells per patient with tdLN NK-like positive (t-test, tumor/blood clones p=0.25, circulating LN-paired p=0.034).^34,38^ i-j. NK-like positive blood T cell average clonal z-score of TEMRA and cytotoxicity signature. k-n. Treatment differences between tumor-blood (t-test, p=0.46) and circulating LN-paired (t-test, p=0.0051) patient clonal average of cytolytic CCR7-CD27-CX3CR1+ signature by Buggert et al. l. Differential expression between ChemoIO versus Naïve tumor-specific blood T cells. m. Feature plot of tdLN UMAP of blood ChemoIO high signature (differentially expressed genes from l. with average log fold change > 0.5 and p adjusted value < 0.05). Linear regression of tumor-specific blood T cell clonal averages of blood ChemoIO high signature versus CCR7-CD27-CX3CR1+ signature by Buggert et al. (R=0.83, p<0.001), blood ChemoIO high signature versus TEMRA signature by Zheng et al. (R=0.87, p<0.0001), and blood ChemoIO high signature versus tdLN NK-like cluster signature (R=0.67, p<0.0001).^34,36^

To assess treatment-induced shifts in clonal differentiation trajectories, we applied the cytotoxic versus dysfunctional classification framework described by Li et al. (**Fig. 5d**).^38^ Compared to treatment-naive uT tumor clones, fewer ChemoIO-treated clones were classified as dysfunctional and more were classified as transitional.^31^ As transitional clones represent a pre-dysfunctional state along the exhaustion trajectory, these findings suggest that neoadjuvant ChemoIO may limit the acquisition of terminal dysfunction and/or partially reinvigorate dysfunctional tumor-infiltrating clones (**Fig. 5d; Supplementary Fig. 6**).

### CD8 T cell clones in or with tdLN lineage respond preferentially to ChemoIO

LN-paired clonal diversity was significantly increased by 46.4% in patients who received neoadjuvant ChemoIO compared to untreated patients (from 58.2 clones to 85.2 clones per 5000 cells), while clonal diversity of clones restricted to the tumor (uT) was not affected by treatment. (**Fig. 5b**) LN-paired clones were also proportionally more expanded within the tumor following treatment. (**Fig. 5b**). Across nearly all transcriptional subtype clusters in both tdLN T cell and tumor T cells, ChemoIO was associated with increased cytotoxicity (Li et al.) and CD39-CD69-stem cell memory programs (Krishna et al) in LN-paired clones (**Supplementary Fig. 5, 6**).^35,38^ *PRDM1* expression, a marker of T cell differentiation, was increased across all tissues and clone types, indicating a global treatment-induced shift in T cell fate (**Supplementary Fig. 7**).^40^ Clonal trajectory analysis further revealed that ChemoIO increased the proportion of tumor LN-paired clones adopting cytotoxic and transitional states, consistent with enhanced functional differentiation following treatment (**Fig. 5d; Supplementary Fig. 6**).

### Circulating LN-paired clones exhibit enhanced cytotoxicity and NK-like features after ChemoIO

The peripheral blood T cell response to ICB therapy particularly relevant in early-stage NSCLC, where systemic surveillance may be required after tumor resection.^48^ Given the inherently cytotoxic profile of circulating LN-paired clones, we next assessed how ChemoIO modulates these subsets across tissues. cLT clones from treated patients exhibited an increased expression of a TEMRA-associated signature (Zheng et al.) in the tdLN, tumor, and blood (**Fig. 5e**).^34^

Within the tdLN’s NK-like cluster, cLT clones from ChemoIO-treated patients exhibited higher cytotoxic and TEMRA scores relative to treatment-naive patients (**Fig. 5f**).^34,38^ This cluster also had a greater cLT clonal diversity in treated patients compared to both untreated patients and treated dLT clones (**Fig. 5f**), indicating that ChemoIO expands the number of clones composed of NK-like T cells within the tdLN. Notably, expression of *KLF2*, *KLF3*, and *NKG7* was increased, even within Naive/CM-like tdLN clusters, suggesting a treatment-induced cytotoxic shift across differentiation states (**Supplementary Fig. 8**).^34,40^

To determine whether the tdLN cytotoxic program extended into circulation, we generated a tdLN-based NK-like signature from differential expression of the tdLN NK-like memory cluster and applied it to peripheral blood clones (**Fig. 5g**). cLT T cells in the blood from ChemoIO-treated patients demonstrated a significant relative expansion of tdLN-based NK-like memory positive T cells (NK-like memory cluster signature z-score > 0) (**Fig. 5h**). ChemoIO also increased the cytotoxic and TEMRA signature expression of NK-like memory-positive cells, mirroring the tdLN clonal treatment response profile (**Fig. 5i, j**).^34,38^

We next aimed to define the transcriptional features of tumor-relevant blood T cells at greater resolution and assess their relationship to the NK-like tdLN-derived response. Differential expression analysis revealed that the peripheral blood T cells of untreated patients were enriched in memory-associated transcription factors such as *TCF7, IL7R, SELL,* and *NELL2* (**Fig. 5l**).^23,34^ In contrast, ChemoIO patients’ blood clones T cells were enriched in NK-like, TEMRA, and effector-related genes *NKG7, FCGR3A, PRF1, IFITM1, IFITM2, IFITM3, GZMB*, etc. (**Fig. 5l**).^34,40,41,45–47^

To determine whether gene expression was biased by any clusters, we defined a ChemoIO circulating blood (CB) response signature (log fold change > 0.25, adjusted p value <0.05) and calculated the patients’ clonal average score within each cluster. Across all clusters, ranging from Naive/CM-like to effector/EM, ChemoIO-treated patients displayed significantly higher scores. The average treatment CB response clonal score was strongly correlated with independent cytotoxicity signatures, including TEMRA (R=0.87) and cytolytic CCR7-CD27-CX3CR1 score (R=0.83) (**Fig. 5m**).^34,46^ Notably, the treatment score was also moderately correlated with our tdLN-based NK-like memory signature (R=0.67) (**Fig. 5n**).

In contrast, bT clones—presumed to originate within the tumor and lacking tdLN lineage—did not exhibit enhanced cytotoxic, NK-like, or TEMRA-like features following ChemoIO (**Fig. 5 k, Supplementary Fig. 6, 8**).^34,38,46^ In addition, unlike circulating LN-paired clones, the NK-like memory-positive populations in bT blood T cells had no difference in proportion, cytotoxicity, or TEMRA signature expression after ChemoIO (**Fig. 5 h-j**).^34,38,39^ Interestingly, cLT clones showed greater clonal expansion after ChemoIO per patient compared to bT clones, but untreated patients’ cLT clones and bT clones had similar levels of clonal expansion (**Supplementary Fig. 8**). These findings highlight a distinct, tdLN-specific effect of ChemoIO that selectively promotes the expansion and activation of cLT CD8 T cells, with minimal impact on bT clones.

Together, our analysis reveals that ICB treatment orchestrates memory-like cell fates locally within the benign tdLN of NSCLC patients and clonal shift in cytotoxicity in clones with tdLN lineage. This is highlighted by the comparison between cLT and bT peripheral blood T cells, where the tdLN-specific TEMRA/NK-like ChemoIO response is only reflected in cLT clones and not bT clones after treatment.

## Discussion

Lung cancer is the leading cause of cancer death in the U.S. and worldwide for both men and women. NSCLC accounts for approximately 80-85% of all cases.^49,50^ Despite advancements in early detection that have increased the population eligible for surgical intervention, up to 55% of early-stage patients develop distant metastatic recurrence within two years after tumor resection and are met with limited therapeutic options and a poor prognosis.^15^

The extent of which tdLNs contribute to systemic anti-tumor immunity is not yet fully understood but carries significant clinical implications. First, the optimal extent of lymph node preservation during lymphadenectomy in resectable NSCLC remains unclear. While this common surgical practice aims to remove regional nodes at risk for metastasis and enable detailed pathological assessment, evidence supporting a survival benefit from systematic lymphadenectomy is lacking.^3,4,6,38,51^ Furthermore, emerging pre-clinical studies suggest that tdLNs may serve as critical sites of T cell activation with the potential to support durable immune surveillance against disseminated disease.^7,16^ Yet, no patient-based studies have directly assessed tumor-specific T cell responses within benign tdLNs and their effects on tumor T cell subset composition.^8,9,53^ This leaves clinical uncertainty regarding the optimal number of lymph nodes that should be resected and therefore a need to determine the baseline role of benign tdLN in the T cell anti-tumor response.^3,4^

Secondly, while phase III clinical trials of ChemoIO have highlighted the promising role of PD-1 ICB therapy in the treatment of early-stage resectable NSCLC, ChemoIO only improves the long-term survival of early-stage NSCLC patients by 10-15%.^11–15^ ChemoIO has been shown to reinvigorate TILs in the TME, however, the systemic protective mechanism of PD-1 ICB has yet to be determined. The clinical implication of this is the timing of treatment in relation to surgical resection (neoadjuvant vs adjuvant).^10^ If tdLN T cells have a response to ChemoIO, neoadjuvant treatment would allow tdLN to support anti-tumor activity prior to their removal. Our study aims to provide insight on the benefits to neoadjuvant treatment by establishing the role of benign tdLN in ChemoIO tumor-relevant T cell response in patients.^10^

In this study, we present the most comprehensive analysis to date of tdLN tumor-specific anti-tumor response in both the number of patient samples and tumor-specific T cell counts, encompassing a control and treatment cohort. We report the largest single cell resolution study of paired tumor and benign tdLN samples. We applied single-cell resolution TCR-based lineage tracing on 672,886 sorted-pure CD8 T cell populations by using the TCR CDR3 region as a molecular barcode to identify and track tumor-relevant clones across tissues.^2,29^ This approach enabled direct identification of 191,471 “tumor-specific” T cells and 2,166 unique clones along with their transcriptional states, with a deep granular analysis of the tdLN, tumor, and circulating blood (n=41). Using this comprehensive dataset, we identified regional lymph nodes as a distinct site harboring tumor-specific CD8 T cells that are responsive to ChemoIO.

The present study is the first to identify two major tdLN-associated transcriptionally distinct clonal subsets: tdLN-tumor TCR-paired clones (dLT) and circulating LN-paired clones (cLT). Through differential expression analysis and literature-based transcriptional signature validations, we found that dLT clones were enriched with *ITGAE, S100A11, CXCR4, PDCD1,* and *TOX*, relating to functions of tissue-residency, activation, and progenitor exhaustion.^21,52^ cLT clones, on the other hand, were enriched in *NKG7, KLRG1, FCRL6,* and *KLF2*, retaining memory cytotoxic potential and migratory signatures consistent with systemic surveillance.^34,42^ Importantly, this transcriptional divergence found in early-memory (naïve/cm-like) tdLN T cells between dLT and cLT clones was continued in the tumor with a differentiated effector-like state. T cells are generally lineage-restricted; therefore, we speculate that the dedifferentiation from effector states to naïve/cm-like states of the 1,689 tdLN clones sequenced is very unlikely.^53^ In addition, within the tumor, tumor-restricted unpaired (uT) clones were transcriptionally distinct from LN-paired clones (increased expression of inhibitor receptors, decreased expression of memory-related genes). Memory-like and cytotoxic E/EM tumor T cell UMAP clusters were significantly contributed by LN-paired clones.

Together, these findings exhibit that the tdLN’s orchestrate a baseline anti-tumor response by transcriptionally differentiating which clones have circulation/surveillance and migration potential (cLT) and which clones have enhanced effector TIL behavior with direct tdLN-tumor lineage (dLT). Furthermore, this tdLN-directed divergence of dLT and cLT impacts the local tumor T cell anti-tumor environment, creating T cell phenotypic memory diversity not attainable by tumor-restricted (uT) clones. These results raise implications for surgical practice. While lymphadenectomy is routinely performed for staging and regional control, these findings indicate it may eliminate a compartment required for systemic immune activation. These results support re-evaluating nodal management in the setting of resectable disease and provide a rationale for preserving benign tdLNs when feasible.

Our findings are the first to demonstrate that ChemoIO directly alters tumor-specific clonality and differentiation trajectories locally in benign tdLN T cell memory state, creating a paramount shift on ChemoIO’s currently known effect beyond exhausted tumor T cell reinvigoration. Specifically, our findings show that ChemoIO selectively enhanced the clonal diversity of LN-paired clones, while unpaired tumor T cell clonal diversity was unchanged. While prior literature has shown ChemoIO to increase clonal diversity, we present that this increase is attributed to tdLN TCR lineage. We hypothesize that the chemotherapy in ChemoIO increases the exposure to new tumor-associated antigens, while the immunotherapy blocking the PD-1/PD-L1 interaction between dendritic cells and T cells in the tdLN promotes the activation of more TCR combinations.

The transcriptional impact of treatment on tdLN tumor-specific T cells was most clearly demonstrated by the tdLN-specific cytotoxic NK-like treatment response of cLT clones. ChemoIO selectively increased the clonal diversity of cLT clones in the NK-like EM cluster within the tdLN. Patients treated with ChemoIO also displayed an increase in cytotoxic/TEMRA-associated signature within the tdLN, while patients naive to treatment remained less activated.^34,38,39^ This treatment response was carried over in circulating blood, where ChemoIO resulted in cLT clones proportionally expanding NK-like T cells and displayed an increase in cytotoxic/TEMRA-associated signatures and genes (*NKG7, FCGR3A, PRF1, IFITM1, IFITM2, IFITM3, GZMB,* etc.).^34,38,39,42^ Remarkably, in tumor-specific blood-tumor T cell clones that had no lineage to the tdLN, there was no difference in cytotoxic/TEMRA function between treated and untreated patients, demonstrating that this response is specific to tdLN TCR clonal lineage.^34,38,39^ These findings indicate that ChemoIO is enhancing the cytotoxic function of cLT clones locally in the tdLN and subsequently also altering the circulating blood LN-paired clonal phenotype. Most importantly, our data provides rationale in favor of neoadjuvant treatment, allowing ChemoIO to harness the tdLN-response prior to resection.

Our work positions the tdLN as a novel target for therapeutic advancements and as a locus of potential immunological mechanisms distinguishing responders from non-responders. In early-stage NSCLC, where the primary tumor is surgically removed, the importance of metastatic surveillance function may be essential and therefore enhancement of cLT clones may prove beneficial for novel therapeutic targets. It will be important to determine if the migratory and cytotoxic memory signatures of cLT clones promote long-term clonal persistence and surveillance for metastatic disease and its relation to patients’ disease-free survival. Future studies should compare the differences within LN-paired clones in tdLN, tumor, and circulating blood between treatment responders and non-responders to identify targetable lymph node-associated programs. Based on our data, future work also could aim to use peripheral blood as a non-invasive metric to determine a patient’s tdLN-specific ChemoIO cytotoxic/TEMRA response. Finally, studies should also evaluate whether the presence or phenotype of these tdLN-linked clones at the time of surgery predicts long-term outcomes.

Limitations to our study include our focus on high-resolution CD8 T cell analysis, which do not describe potential contributions of other immune populations within the tdLN, including CD4 T cells, antigen presenting cells, and components of the innate immune system. Future work should aim to establish how other populations within the tdLN may support the tdLN T cell anti-tumor response. In addition, while an extensive effort was put forth to provide QC and validation towards our stringent “tumor-specific” T cell categorization, limitations of this study include the absence of known tumor antigens, which precludes direct identification of tumor-specific T cells.

In conclusion, we define the benign tdLN as a clinically relevant site of CD8 T cell engagement in early-stage NSCLC. tdLN-derived clones give rise to transcriptionally distinct effector populations, which are further modified by PD-1 blockade. Their presence in circulation may represent a reservoir of persistent antitumor immunity. These findings provide a foundational shift and a rationale for preserving and therapeutically targeting tdLNs in the treatment of resectable NSCLC.

## Methods

### Clinical sample collection, imaging, and pathology evaluation

Tumor, tumor-draining lymph node, and peripheral blood samples were collected under an Institutional Review Board–approved protocol (#23-04025976) at Weill Cornell Medicine, under the study titled Mechanism of T-cell memory differentiation in a lung cancer model and the role of PD-1 blockade on the tumor draining lymph nodes of NSCLC patients. Samples were obtained prospectively through the Thoracic Surgery Biobank, with informed consent. Pre-treatment and post-treatment CT imaging was reviewed independently by board-certified pulmonary radiologists to assess radiographic features and treatment response. Resected tumor and lymph node specimens were processed and reviewed by a board-certified pulmonary pathologist to confirm histologic diagnosis and treatment-associated changes.

### Clinical matching and sample selection process

As the study focused on patients with early-stage disease, the study only included patients with stage I-IIIa cancer at the time of resection. Patients not treated with ChemoIO were to have no prior NSCLC treatment. One patient, bb1927, had metastatic disease found in the lymph nodes after surgical resection, pathologic assessment, and was also considered to have oligometastatic disease. We collected tumor mutations from each patient stored within the biobank. Patients with targetable mutations (EGFR and ALK1) were excluded from this study.^15^ Blood, tumor, and regional benign lymph node tissue samples were collected from all patients within the study. Patients were also clinically matched for smoking status, gender, stage, and residual viable tumor in ChemoIO-treated patients.

### Flow cytometry and cell sorting

Prior to sequencing, immune cells in single cell suspension were thawed and stained for sorting and were sorted using the flow cytometry for CD45 CD3 and CD8.

### scRNASeq and scTCRSeq and demultiplexing and read processing

To align raw single cell RNA Sequencing and single cell V(D)J sequencing data we used Cell Ranger version 8.0 using the *multi* program. The GEX GRCh38-2024-A human reference genome was used for transcriptional reference along with VDJ GRCh38-2024-A, human reference for V(D)J. Aligned transcriptional data is processed using Read10X using R version 4.4.1. Seurat version 5 was used to create a Seurat object with the following parameters: min.cells set to 3 and min.features parameter set to 200.^55^ Patient’s where the overall number of features were less than 200 per cell were removed. TCR V(D)J data was then processed by combining the filtered_contig_annotations.csv and clonotypes.csv dataset to get the CDR3 alpha and beta amino acid sequence for each cell, which was then added as metadata to the Seurat object. If T cells were not sequenced for both TCR and RNA sequencing, they were removed from the dataset. T cells that did not have both beta and alpha chains of the TCR sequence were removed.^1,2,31,34,35^

### Defining T cell TCR clones and tumor-specific clones

T cells were in the same clone if they shared the same TCR alpha and beta genes. T cell clone size was defined as the number of cells within the given shared clone. Clonal diversity refers to the number of unique TCRs in a given population (ex: patient, or cluster per patient). Tumor-specific clones were expanded (clone size >=10), non-viral T cell clones that have TCR lineage with the tumor are tumor-specific. Viral clones were defined as clones with TRC alpha and beta chains found on the human viral VDJ database (https://vdjdb.cdr3.net/), which lists TCR CDR3 amino acid sequences that relate to known viral antigens.^33^

### scRNA-seq data integration and clustering

Our T cell single RNA sequencing database required thoughtful integration due to the large number of patient sample cells sequenced for one cell type (CD8 T cells). Due to the different tissue types, treatment, and treatment response, we aimed to minimize over-correction. We tested 4 different integration methods (no integration, RPCA, CCA, and harmony) as well as different anchor values for RPCA and CCA methodology. We integrated based on patient ID. When no integration method was applied, each cluster was predominantly defined by patient ID, displaying the need for integration. RPCA integration, using an anchor value of 15, ultimately was the best integration method. Patients with the same treatment (either Naïve to treatment or ChemoIO) were clustered in a similar pattern. In addition, T cells are clustered into biologically relevant T cell subsets. Each UMAP subset (ex: tumor-specific tumor T cells, tumor-specific tdLN T cells, etc.) was re-normalized and re-integrated. Each UMAP had its own PCA analysis, which determined the number of PCAs required for dimensional reduction. Dimensional reductions were performed as instructed by Seurat.^,55^ T cell cluster subsets were then defined by the up or down regulation of memory markers (ex*. SELL, LEF1, TCF7*, etc.), effector-related genes (ex*. CD44, PRF1, GZMB, EOMES*, etc.), dysfunctional/inhibitory receptor genes (ex. *TOX, CTLA4, HAVCR2*, etc.), and proliferation markers (ex, *MKI67*).^1,2,31,32,34,35,38–42^ In addition, the *FindAllMarkers* function was used to determine which genes were differentially expressed within each cluster. ^55^ These results were used to determine both T cell cluster function, and if further dimensional reduction were required. Cluster subset labels were then validated using modular scores of T cell signatures, which received z-score normalization, found in the literature.

### Gene signature scoring analysis and statistical tests

To compare phenotypic differences between groups of T cells (ex, between clusters, between treatments, between tissues, etc), a modular score was made of signatures from the literature using *AddModuleScore*.^55^ These scores were then normalized within a UMAP via z-score normalization (mean of zero, standard deviation of one). Three forms of analysis were made: comparing individual T cell scores, averaging the score across all cells within the same clone, and averaging each clonal average per patient. While comparing T cell score averages give single-cell level insight, averaging across clones provides information regarding a clonal state. In addition, due to the large number of T cells, a small score difference can lead to a statistical difference. Therefore, using clonal averages decreases the total number of points on the statistical test, making the statistical significance more powerful. When comparing two groups, a test was performed. A paired T test was performed if changes were compared within the same group (i.e. the same clone in different states, or comparing the same patient in different states or clonal pairing). Because bb2050b did not have blood samples collected, they were removed from the paired t test comparing cLT and dLT. Similarly, when comparing paired phenotypic changes between tumor-blood and cLT clones per patient, patient bb2054 was removed due to not having any tumor-blood clones. When comparing multiple t tests within the same dataset, an ANOVA test was also performed to reduce type 1 error.

### Relative clonal diversity and clone pairing expansion analysis

The number of T cells sequenced for each patient sample ranged from a few thousand to 20,000 cells. Because of this, the absolute number of tumor-specific T cells for each tissue sample also ranges from a few thousand to 13,000. Therefore, to compare the clonal diversity between patients, we normalized the number of clones found by the number of cells within the clonal pairing. We followed this by multiplying the normalized clonal diversity by 5000 to get the number of unique clones per 5000 tumor-specific cells.

To compare the relative proportion of clonal pairing (ex: unpaired tumor only versus LN-paired clones) per cluster, we calculated the proportion of paired clones within a given cluster and divided it by the patient’s specific overall clonal pairing portion. Therefore, a relative proportion 1 for each clonal pairing represented equal proportions within a cluster as the patient’s proportion.

### Cluster clonotype abundance index

The relative abundance of a clonotype pairing (ex: uT, dLT, cLT, bT) within a UMAP cluster describes whether a given clonotype is abundant within a cluster compared to its general population proportion within a tissue for a given patient. The relative abundance index of a clonotype within a given cluster is calculated by dividing the cluster’s clonotype T cell proportion by the patient’s global proportion of that clonotype within the tissue. A cluster clonotype abundance index of 1 would indicate that the cluster is not dominated by any clonotype. If all clusters and clonotypes have an abundance index of 1, it would indicate that clonotypes and TCR tissue lineage do not influence UMAP clustering and clonotypes do not have preferential T cell subtypes. If a clonotype within a cluster has an abundance index greater than 1, then the clonotype is abundant and dominant in the cluster. This means that the cluster’s transcriptional profile and dimensional reduction are derived and dominated by the given clonotype and tissue TCR lineage; consequently, tissue TCR lineage clonotypes do have preferential T cell subtypes within a tissue’s UMAP.

## Acknowledgments

We thank Julissa Murillo, Abu Nasar, and Arshdeep Singh for clinical support. Dr. Jenny Xiang of the Genomics Resources Core Facility for professional advice. This work was supported in part by LUNGevity, The Lung Cancer Foundation of America, NIH grant 3UH3CA244697-04S1, The Neuberger Berman Foundation Lung Cancer Research Center and support from the Meyer Cancer Center, NSF Graduate Research Fellowship Program (GRFP) awarded to R.H. (DGE-2139899). The funding organizations played no role in experimental design, data analysis or manuscript preparation.

## Author contributions

J.V.V and O.E. supervised this study. R.H. and J.V.V designed the experiments. R.H. performed the computational analysis and wrote the manuscript. T.D.C and S.T. prepared clinical tissue samples for sequencing. R.H., S.T. and O.D. performed biobanking. L.V. provided methodological and computational guidance. J.V.V. edited the manuscript. G.E., M.M., N.K.A. gave suggestions. A.S. coordinated the clinical sample acquisition.

