## Supplement Figures for "Tumor-specific draining lymph node CD8 T cells orchestrate an anti-tumor response to neoadjuvant PD-1 immune checkpoint blockade"

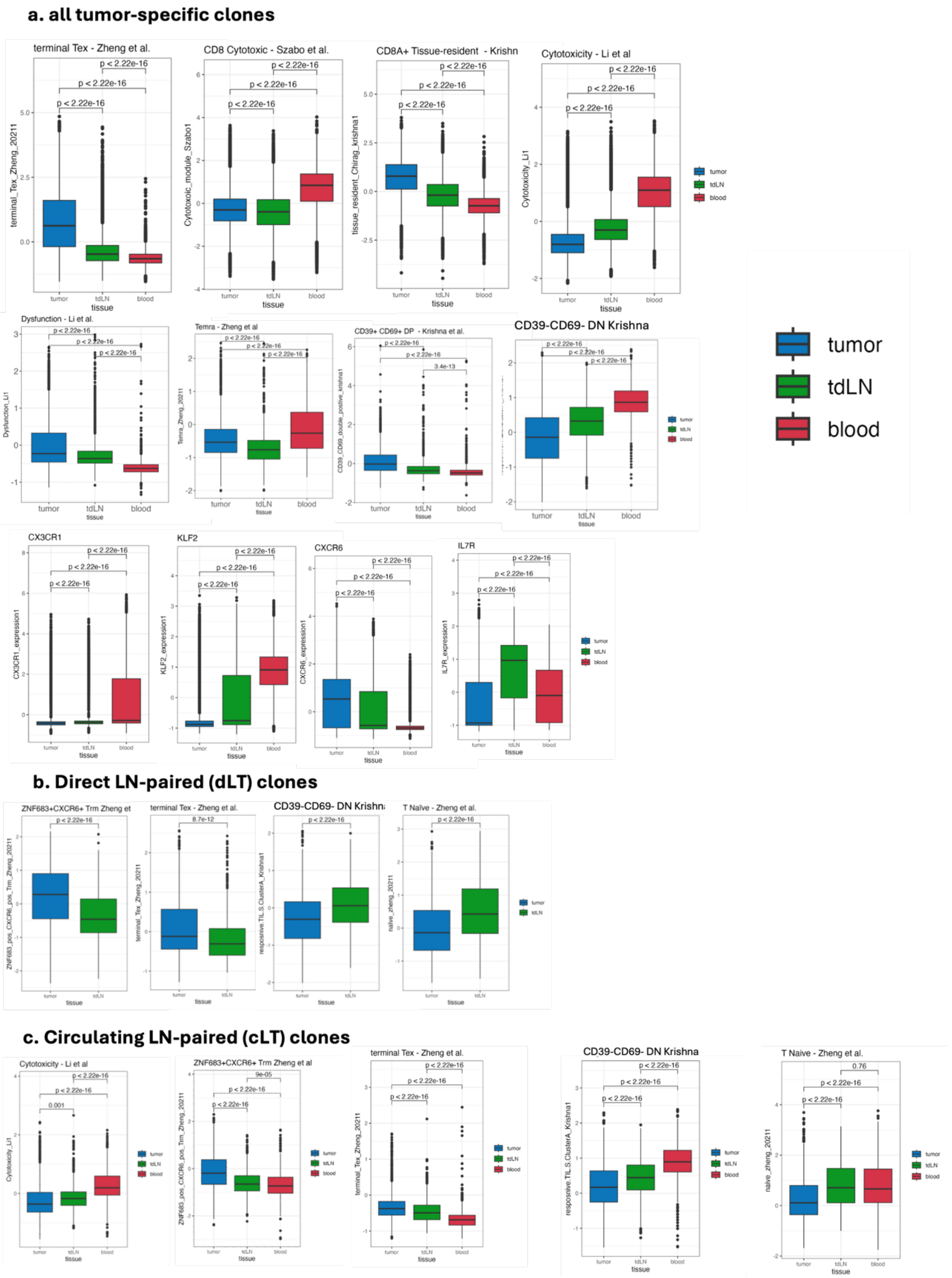


*Supplement Figure 1: Tumor-specific T cell function within the tumor, tdLN and blood. a. Clonal average z-score within each tissue for tumor-specific clones (t-tests and ANOVA). b. Clonal average z-score within the tumor and tdLN for all direct LN-paired (dLT) clones (paired t-test). c. Clonal average z-score for each tissue for all circulating LN-paired (cLT) clones (t-test and ANOVA).*

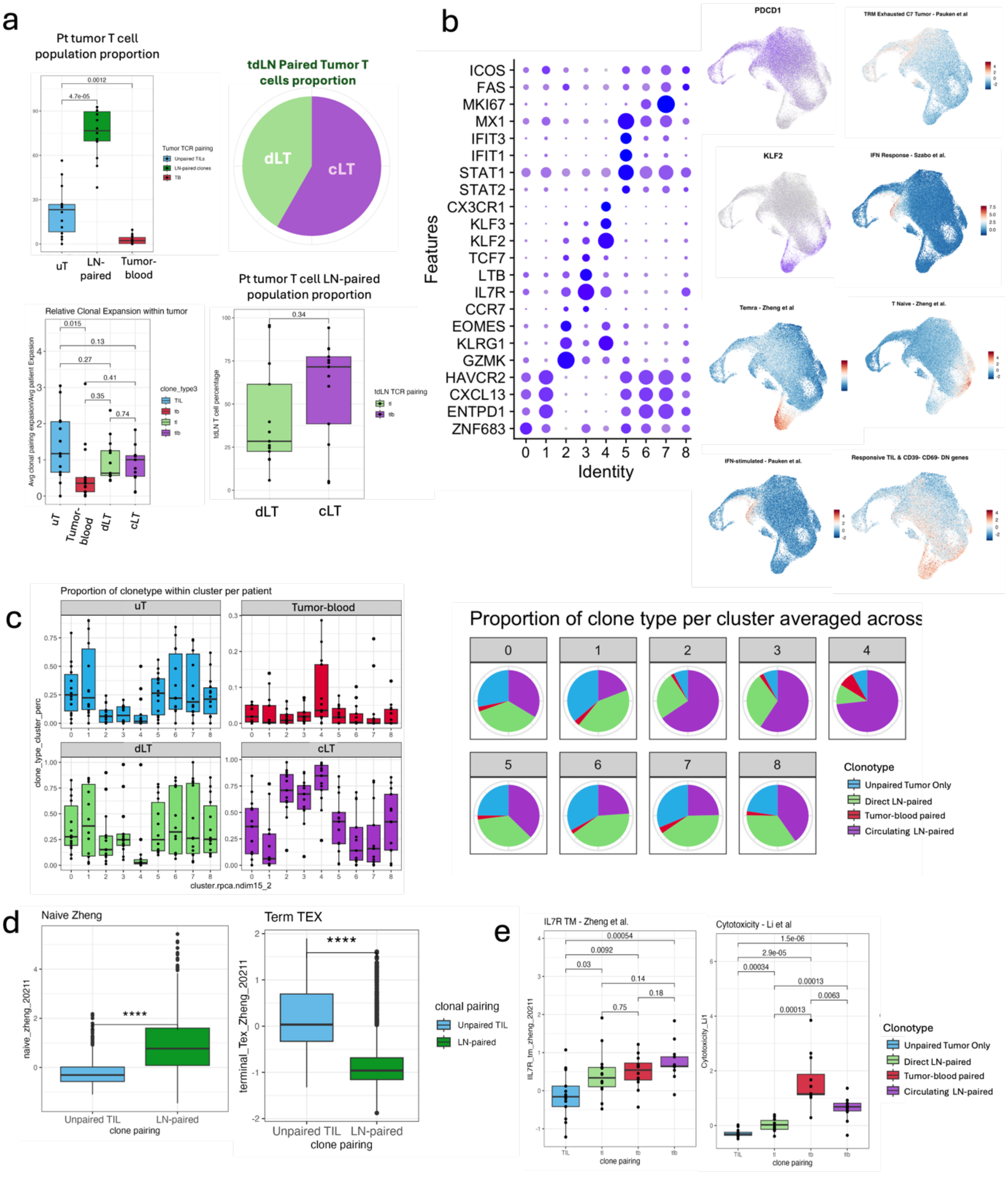

*Supplement Figure 2: Tumor-specific T cell function within the tumor, tdLN and blood. a. Analysis of clonal population proportion and relative clonal expansion analysis between each clonotype. b. UMAP cluster analysis. Left a DotPlot of highly expressed genes in each cluster. Right supplemental FeaturePlots of genes and signatures. c. Proportion of each clonotype within each cluster. d. Clonal average signature expressions between unpaired tumor only clones and LN-paired clones. e. Supplemental clonal average signature expressions between all clonal types.*

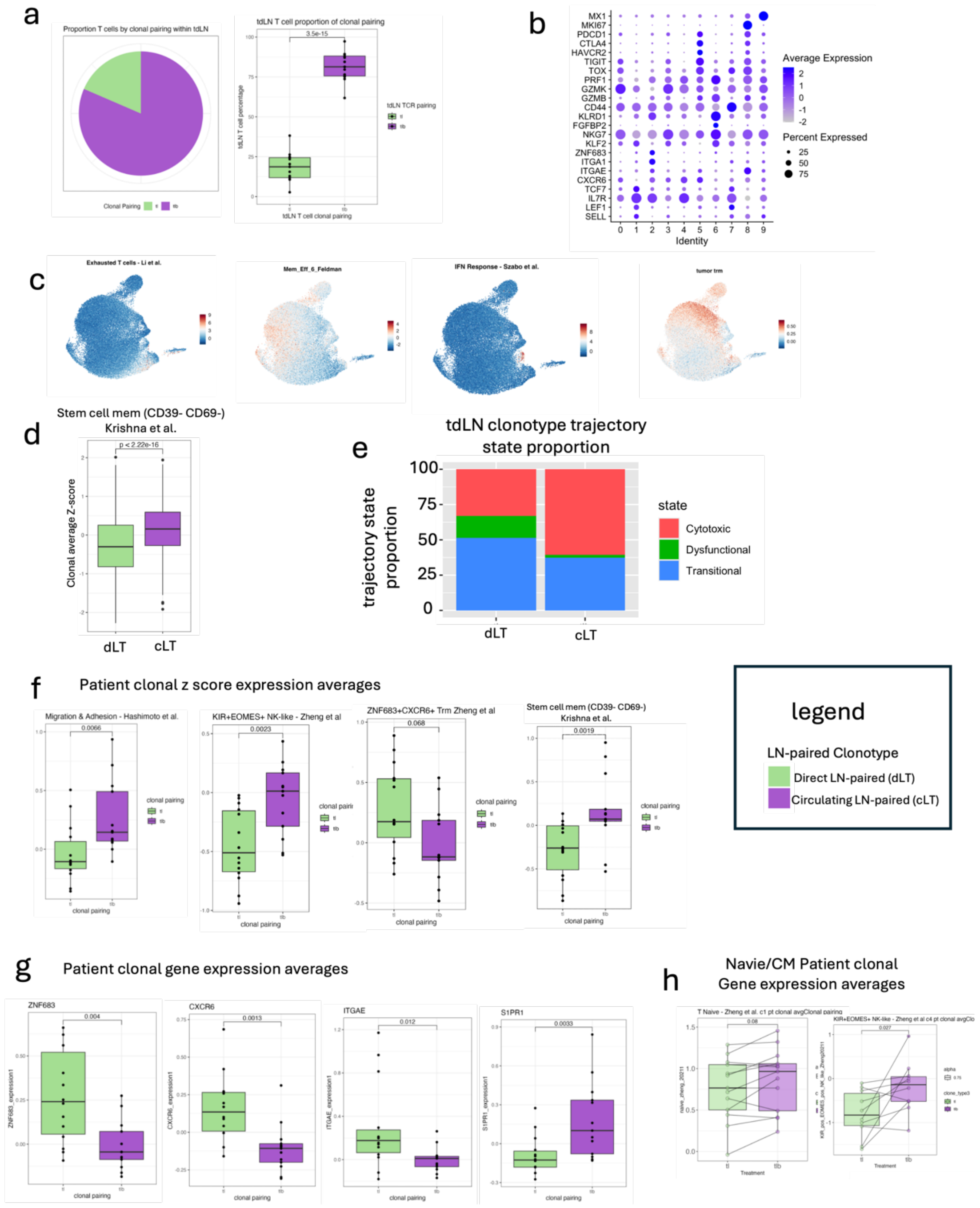


*Supplement Figure 3: Two distinct LN-paired clonal memory phenotypes found within the tdLN. a. TdLN T cell proportion by clonal type per patient (paired t-test). b. DotPlot of highly expressed genes in each cluster. c. Supplemental FeaturePlots of genes and signatures. d. Clonal average z-score stem cell memory signature (Krishna et al., t-test). e. TdLN clonotype trajectory state proportion for dLT and cLT clones. f. Supplemental patient clonal average z-score expression of signatures between clonotypes within the tdLN (paired t-test). g. Supplemental patient clonal average z-score expression of genes between clonotypes within the tdLN (paired t-test). h. Patient clonal average z-score within the Naïve/CM tdLN UMAP cluster (paired t-test).*

a

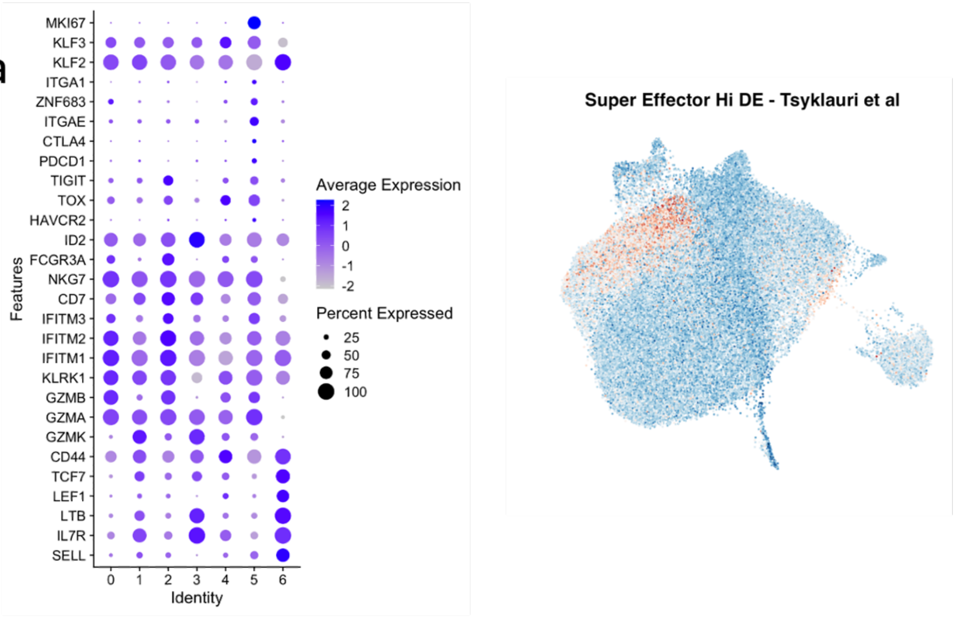


b

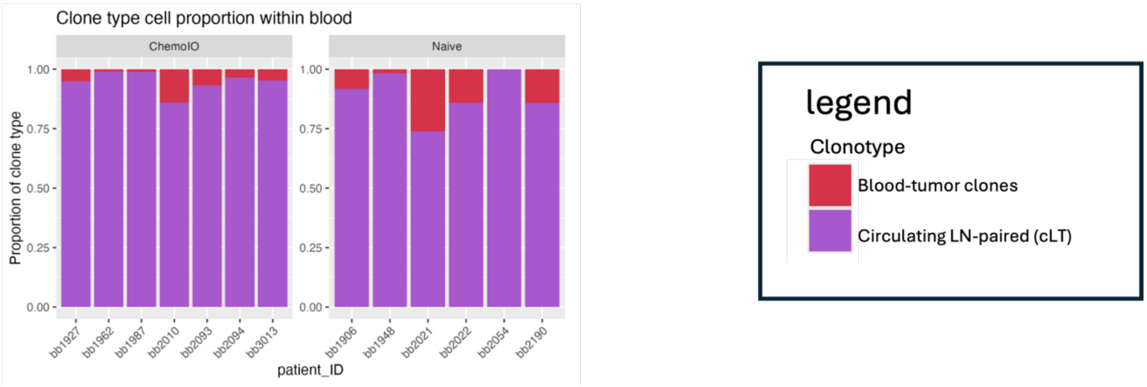


*Supplement Figure 4: Contribution of LN-paired clones in tumor-specific circulating blood T cells. a. Left DotPlot of highly expressed genes in each cluster. Right supplemental FeaturePlot. b. Clonotype proportion per patient grouped by treatment cohort.*

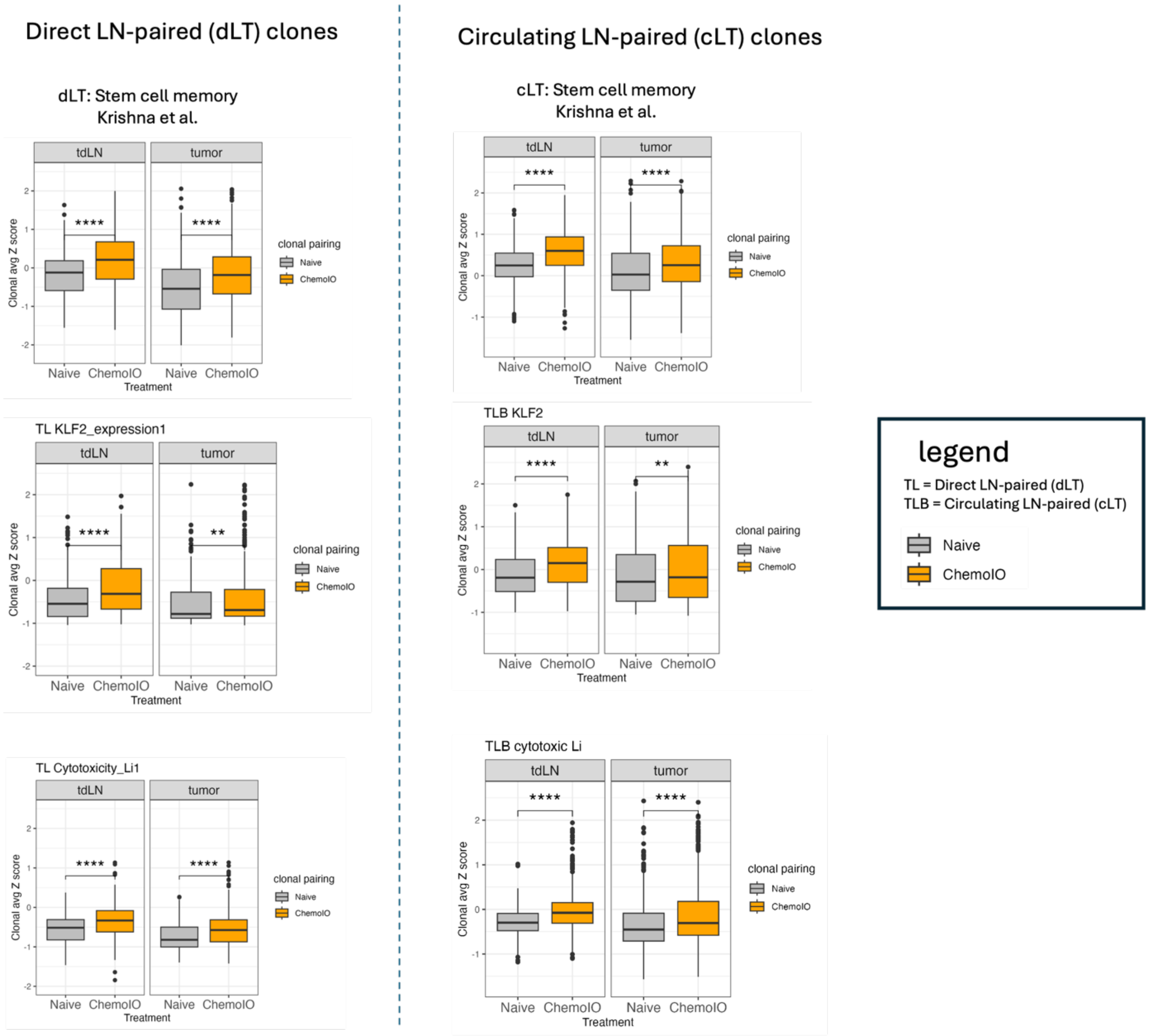


*Supplement Figure 5: Comparison of ChemoIO vs Naïve treated patients' direct (left) and circulating (right) LN-paired clonal average z-score differences between the tdLN and tumor (t-test).*

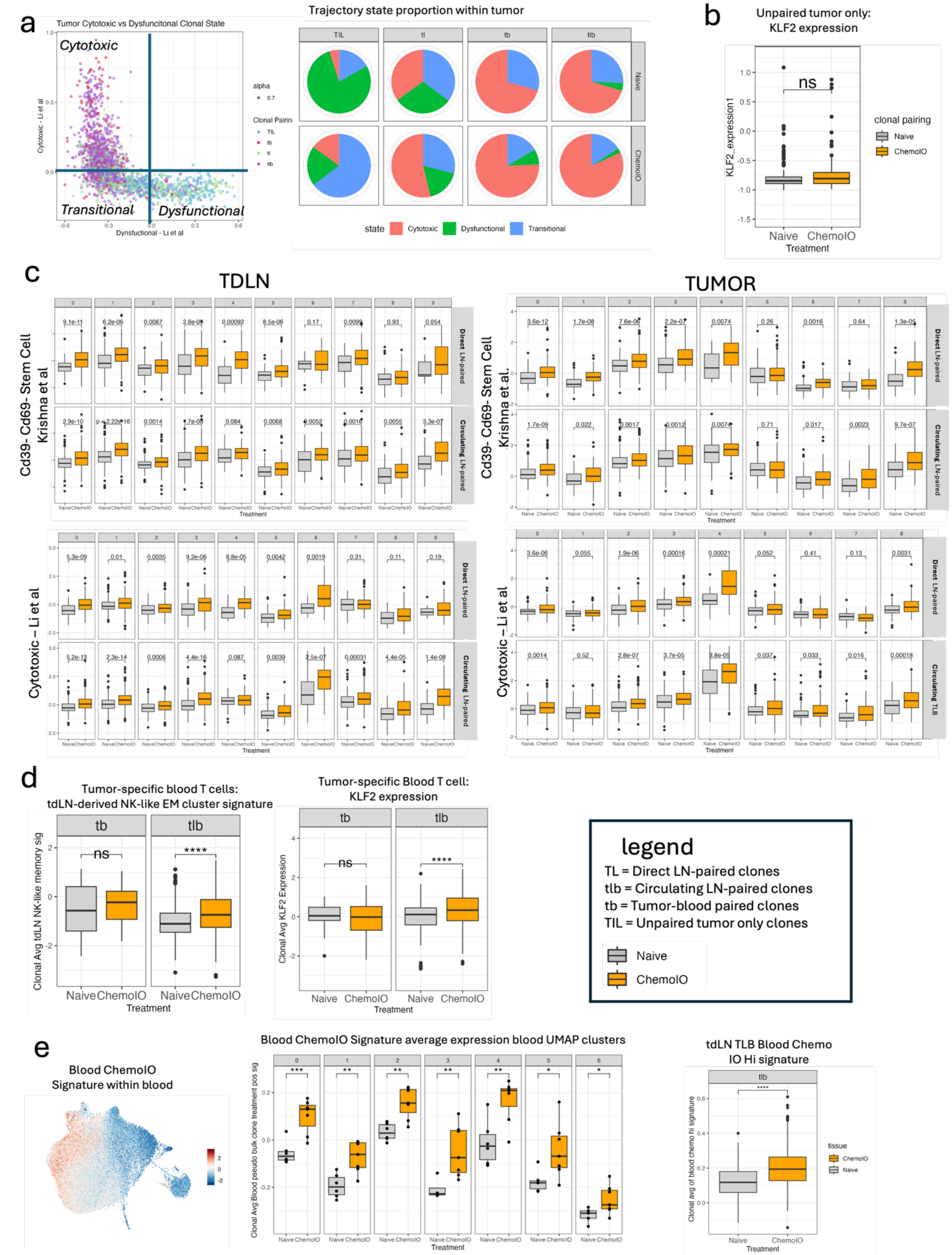


*Supplement Figure 6: ChemoIO response in tdLN and tumor. The quantification of cytotoxic vs dysfunctional categorized clones. Supplementary plots regarding ChemoIO blood signature. a.*

*Clonal trajectory cytotoxic versus dysfunctional state analysis within the tumor. b. Unpaired clonal expression of KLF2 within the tumor between treated and untreated patient (t-test). c. Clonal average z-score of CD39- CD69- stem cell memory (top) and cytotoxic (bottom) per each cluster and LN-paired clonotype within tdLN and tumor between treated and untreated patients' clones (t-test). d. Blood T cell clonal average z-score between treated and untreated by clonotype (t-test). e. Left ChemoIO tumor-specific blood signature FeaturePlot on blood UMAP. Middle signature applied to each blood UMAP cluster (patient clonal average) by treatment cohort. Right signature applied to the tdLN cLT clones between treated and untreated (t-test).*

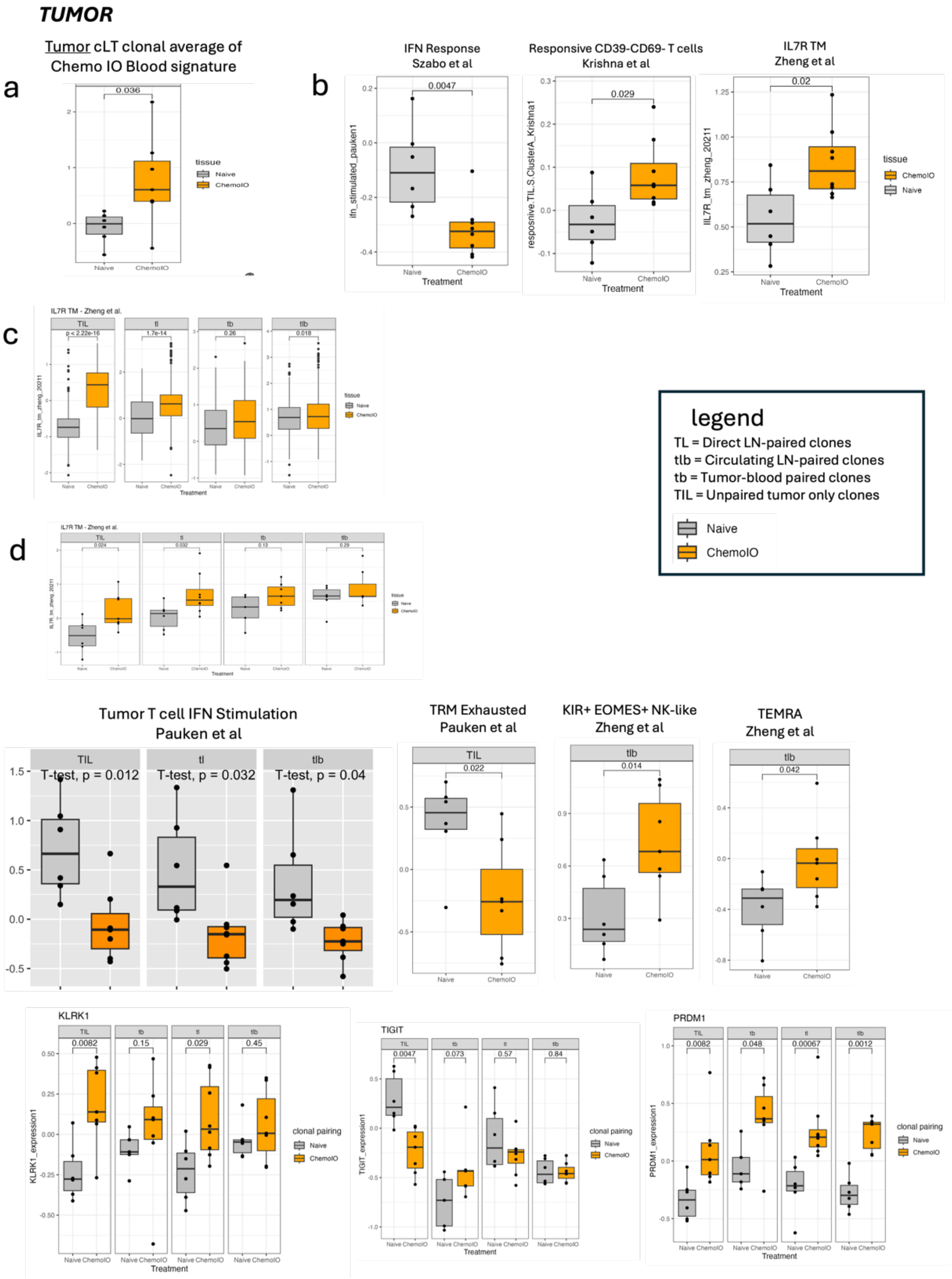


*Supplement Figure 7: ChemoIO response in unpaired tumor-only clones and LN-paired clones within the tumor. a. Tumor cLT clonal average ChemoIO blood signature patient clonal average z-score by treatment cohort (t-test). b. Patient clonal average z-score of all tumor T cells of the signature (t-test). c. Clonotype-specific clonal average z-score IL7R TM signature expression by treatment cohort in the tumor (t-test). d. Patient clonal average z-score by treatment per clonotype within the tumor for supplemental signature and gene expressions (t-test).*

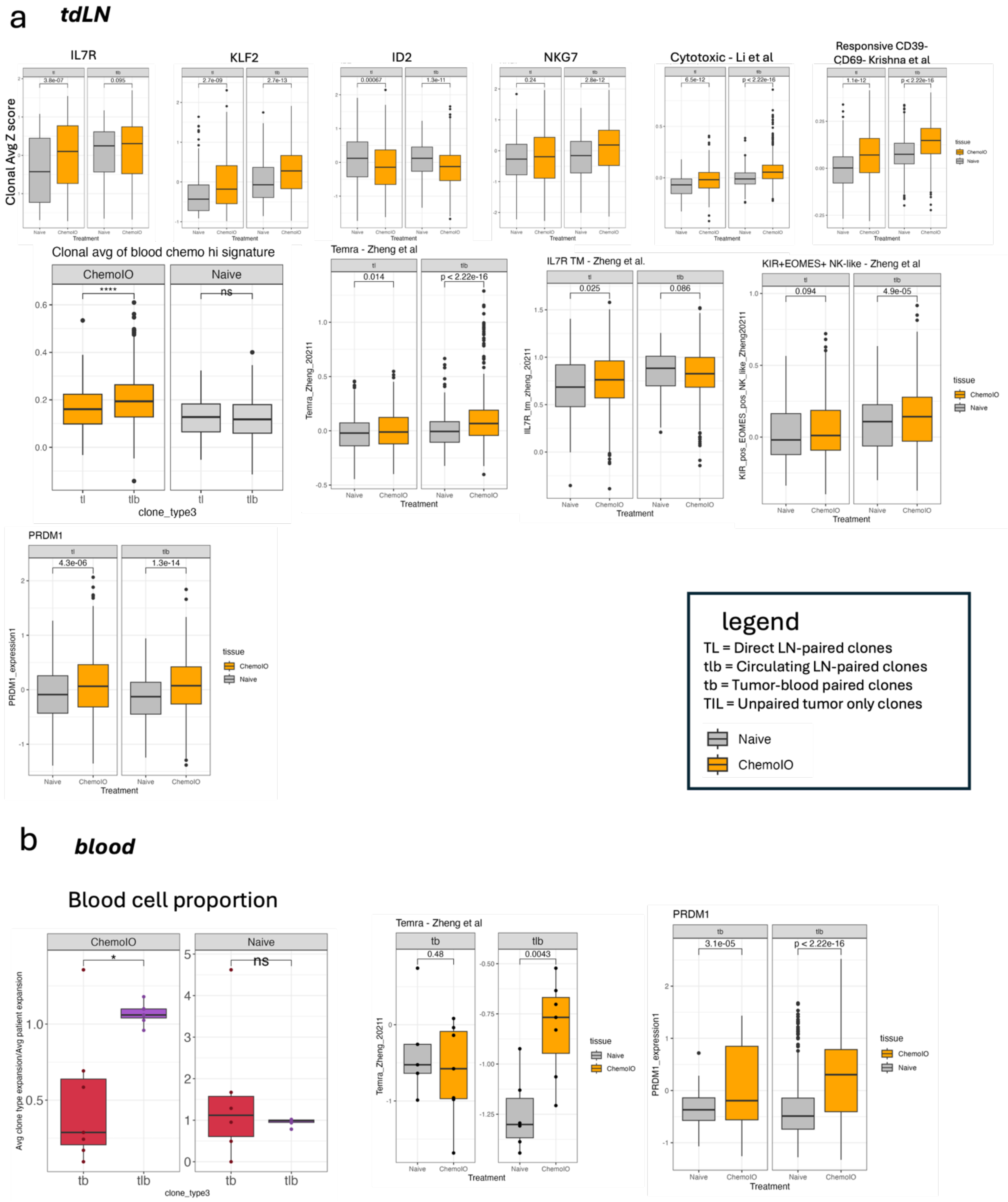


*Supplement Figure 8: ChemoIO response in tdLN and blood LN-paired clones within the tumor. a. Clonal average z-score of supplemental signatures and genes within the tdLN by treatment and clonotype (t-test). b. Treatment differences within the blood. Left proportional clonal expansion of clonotypes per patient (paired t-test). Right supplemental signature and genes relating to treatment changes within the blood per clonotype.*
